# Intravenous BCG Elicits a Sustained Myeloid IFNG Response Signature Associated with Early TB Control

**DOI:** 10.64898/2026.11.25.768954

**Authors:** James D. Dahlvang, Paul L. Maurizio, Molly A. Robertson, Allison N. Bucsan, Chelsea C. Lehman, Farida Laboune, Amy R. Henry, Laura E. Via, April M. Walker, David R. Ambrozak, Melanie Cohen, Julie Laux, Iyadh Douagi, Taylor Farley, Justin L. Gaines, Mohau S. Makatsa, Chetan Seshadri, Forrest Hopkins, Michael C. Chao, Sarah M. Fortune, JoAnne L. Flynn, Daniel C. Douek, Mario Roederer, Patricia A. Darrah, Robert A. Seder

## Abstract

Bacille Calmette-Guérin (BCG) administered intravenously (IV) to rhesus macaques can prevent pulmonary tuberculosis (TB) infection. To establish transcriptional correlates of protection, we generated longitudinal CITE-Seq data across blood and lung after IV BCG in macaques and compared it to clinically approved intradermal (ID) BCG. We define an interferon gamma (IFNG) response signature in alveolar macrophages (AMs) comprising 61 genes with diverging expression between IV and ID BCG from 4 to 16 weeks after vaccination that was also enriched in AMs early after TB challenge. The signature was recapitulated in blood monocytes (IV vs. ID) and associated with protection in an independent cohort of IV BCG-vaccinated macaques. Cell-cell communication analysis inferred increased ligand-receptor interactions in the airway after IV BCG, including those between multiple IFNG-producing T cell subsets and AMs. The sustained IFNG response signature in AMs after IV BCG provides a new correlate and potential mechanism of early TB control informing future vaccine design.

## INTRODUCTION

Tuberculosis (TB) remains the leading cause of mortality attributable to a single infectious disease worldwide, accounting for more than one million deaths in 2024^1^. The only licensed vaccine for TB is live-attenuated Bacille Calmette-Guerin (BCG), which is administered intradermally (ID) at birth in TB-endemic countries. ID BCG limits disseminated TB in infants and young children but fails to provide durable protection against pulmonary TB through adolescence and adulthood, allowing continued transmission^2–6^. Thus, developing vaccines that mediate improved protection against pulmonary TB is a primary goal for limiting the spread of TB disease. TB vaccine development is hindered by the lack of clearly defined immune correlates of protection in humans or pre-clinical animal models. In recent years, high-level protection against TB in macaques has been achieved after live-attenuated whole organism vaccine approaches including mucosal BCG^7,8^, aerosol (AE) MtbΔSigH^9,10^, and intravenous (IV) BCG^11,12^, as well as with intramuscular RhCMV/TB^13^. Across several studies in a highly pathogenic rhesus macaque model, IV BCG immunization prevented TB infection and IFNG release assay (IGRA) conversion in >50% of macaques challenged with Mtb Erdman, suggestive of very early control^11,12,14^. In contrast, ID BCG at the approved human dose confers a ∼1-log reduction in Mtb burden compared to unimmunized infected macaques (∼10^6^ Mtb) during a similar 12-week challenge phase^11^. These data provide a framework to compare BCG delivered by the IV route that provides optimal protection by preventing infection, with ID BCG as it is currently delivered across the world to inform next generation vaccine development.

IV BCG immunization elicits robust cellular and humoral immunity in the periphery and in the airway (bronchoalveolar lavage, BAL). Multivariate immune analyses based largely on flow cytometry and systems serology data identified the number of polyfunctional, Th1/Th17 cytokine-producing CD4 T cells and NK cells in the BAL, as well as complement-fixing IgM and NK-cell activating antibody in plasma as correlates of protection^12,15^. To substantiate adaptive immune mechanisms of protection, *in vivo* depletion studies were performed in macaques. Importantly, depletion of B cells in macaques during IV BCG immunization had no effect on protection against TB^16^. In contrast, depletion of CD4 T cells or CD8ɑ-expressing cells prior to Mtb challenge in IV BCG-immunized macaques abrogated protection, demonstrating that T cells are necessary for protection^14^. Of note, although CD4-and CD8ɑ-depletion in IV BCG-immunized macaques resulted in high Mtb bacterial burdens over the course of infection, they retained the ability to restrict the number of Mtb bacilli that established infection, as determined by Mtb barcode analysis. These data suggest a potential role for innate immunity in early TB control after IV BCG. In mice, IV BCG can reprogram the myeloid compartment in bone marrow or lung in an IFNG-dependent manner, resulting in heightened innate recognition of homologous (or heterologous) pathogens by monocytes and macrophages in the lung^17–21^. As alveolar macrophages (AMs) are the first host cells to encounter Mtb infection^22,23^, their ability to respond quickly to infection may be critical to initial control. This potential mechanism for mediating early control of TB in macaques following vaccination has not been fully elucidated.

Single-cell RNA sequencing (scRNA-Seq) provides a sensitive and unbiased approach to profile cell type-specific immune responses and cell-cell communication that may underlie protective correlates. In macaque models, scRNA-Seq has been applied to permissive versus restrictive granulomas and active versus latent lung tissue after primary Mtb infection or reinfection, revealing immune responses associated with TB resolution^24–27^. In the context of vaccination, we and others analyzed limited BAL transcriptional datasets from macaques after immunization with ID or IV BCG, identifying enhanced Th1/Th17 T cell signatures and CD4 T cell–macrophage intercellular communication associated with IV BCG; however, without Mtb challenge outcome, correlates of protection could only be inferred^11,28^ A transcriptional analysis several weeks after challenge with a less pathogenic Mtb strain (CDC1551) in AE MtbΔSigH-immunized cynomolgus macaques induced a broad and balanced Type I and Type II IFN responses in the BAL compared to AE BCG and was associated with improved outcome in this more resistant model^10^. Prior bulk transcriptional analysis of blood from macaques after IV BCG revealed an early (d2) innate Type I IFN response signature that predicted subsequent T cell responses in BAL and protection^29^; however, bulk transcriptional profiling lacks the depth to characterize cell-type specific gene modulation in response to vaccination or infection.

Here, to define immune signatures of protection, we generated a longitudinal CITE-Seq (Cellular Indexing of Transcripts and Epitopes by Sequencing^30^) dataset before and after IV and ID BCG vaccination and early after TB challenge from blood, BAL, and lung tissue to characterize the cell-type specific features and interactions that lead to the profound differences in early and sustained protection against TB. The data reported here show broad T cell-derived *IFNG* gene expression and sustained IFNG responsiveness in myeloid cells after IV BCG. We define a 61-gene IFNG response signature with temporally divergent expression in BAL AMs and blood monocytes from IV vs. ID BCG-immunized macaques after vaccination as well as in AMs 11 days post-challenge. In an orthogonal dataset generated from an independent cohort of IV BCG immunized macaques, the 61-gene signature was enriched in blood monocytes from animals with greater protection against TB disease. These findings suggest the sustained IFNG responsiveness in blood monocytes and BAL AM after vaccination and infection may be signatures of a highly protective response and facilitate early TB control in the airway.

## RESULTS

### Generation of a multi-tissue CITE-Seq atlas of BCG vaccination in rhesus macaques

Eighteen rhesus macaques were vaccinated with either a standard human dose of ID BCG (∼5 × 10^5^ CFU, n = 9) or IV BCG (∼5 × 10^7^ CFU, n = 9) and a subset of each group (n = 3) was challenged with Mtb (Erdman) 22 weeks later (**Figure 1A** and **Table S1**). PBMC and bronchoalveolar lavage (BAL) samples were cryopreserved before (Pre) and after (Wk4 and Wk16) BCG vaccination; cells from BAL and lung tissue were cryopreserved 11 days (D11) after Mtb challenge. Both IV and ID BCG vaccinated animals generated antigen-specific T cell responses, but IV BCG animals exhibited a substantial influx of T cells into the BAL and larger CD4 T cell cytokine responses to Mtb whole cell lysate (WCL) stimulation in BAL and PBMCs (**Figure S1A-C**). These data indicate ID and IV BCG immunization in this study recapitulate immune responses reported in prior efficacy studies^11,12^.

**Figure 1.**
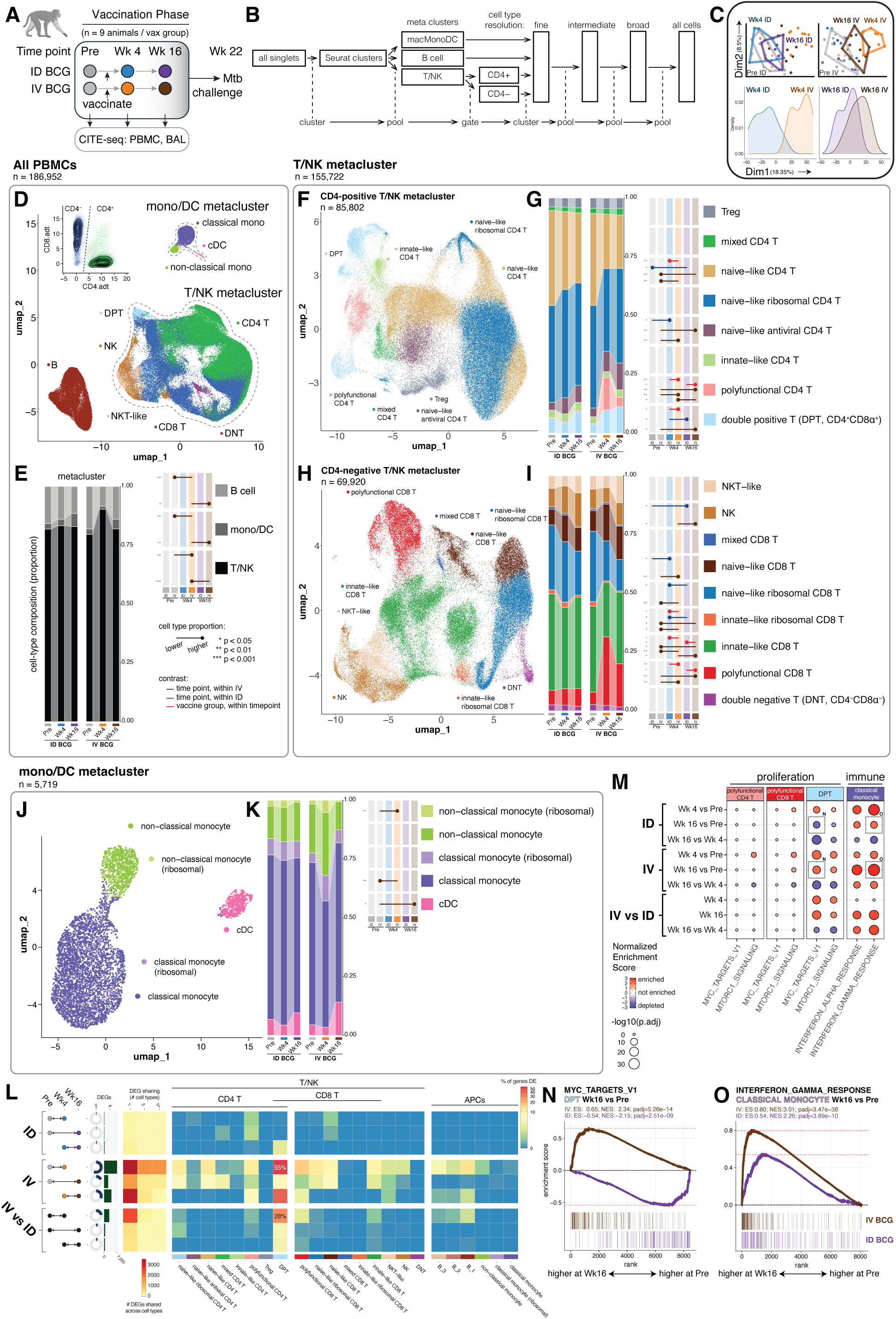
Study design and single-cell CITE-Seq analysis of vaccination phase PBMCs. **(A**) Study design and color coding for macaques vaccinated with ID BCG (n=9) or IV BCG (n=9). PBMCs and BAL were collected before (Pre) and after (Wk4, Wk16) BCG. **(B)** Overview of cell type clustering, counts, and annotation workflow; see **Figure S2A** for extended detail. **(C)** Multidimensional scaling (MDS) of PBMC data pseudobulked by sample at each timepoint (top). Colored polygons include samples from macaques within a timepoint vaccine group, ID BCG (top-left) and IV BCG (top-right). Ridgeline plots (bottom) represent distribution of Dim1 values for ID and IV BCG at Wk4 (left) and Wk16 (right). **(D)** Uniform Manifold Approximation and Projection (UMAP) embedding of scRNA-Seq profiles of vaccination phase PBMCs (*all cells*) with ADT partitioning of CD4-positive and CD4-negative metaclusters (inset). **(E)** Cell type composition (metacluster proportion of all cells) within ID and IV BCG samples at each timepoint (stacked bars, left). Significant differences (*P*.adj < 0.05, BH FDR) between timepoints within IV (brown) or ID (blue), or between vaccine groups within timepoint (red) determined by generalized linear modeling are depicted with color-coded directional lollipops. (**F**-**K**) UMAP of PBMCs within the CD4-positive T/NK metacluster (**F**), CD4-negative T/NK metacluster (**H**) and mono/DC metacluster (**J**). Cell type composition and significant changes, as in (**E**) for each metacluster (**G**, **I**, **K**). **(L)** Differential expression (DE) analysis of genes in PBMCs for each timepoint contrast within ID (top) or IV (middle), or between vaccine groups across timepoints (bottom); contrasts are labeled at far left. Percent (donuts) and number (bars) of DEGs (of all genes profiled) across all cell types and DEG sharing heatmap depicting the number of DEGs (color scale below) shared across 1, 2, or 3 (or more) cell types. Within the T/NK cells and APCs (B cells, mono/DCs), heatmap illustrating the percent of DEGs (color scale on right) within each cell type (labeled below) for each contrast. Significant differential expression was defined as FDR < 0.1 (BH). (M) Selected GSEA results. Normalized enrichment scores (NES) and associated *P* values based on GSEA of rank-ordered signed *P* value lists (−log10*P* × sign(logFC)) derived from DE analysis. Shown are selected enriched Hallmark pathways (below) within polyfunctional CD4 T, polyfunctional CD8 T, DPT cells, and classical monocytes for each contrast (left). Complete GSEA is presented in **Figure S5**. (**N**-**O**) Barcode plots of MYC Targets v1 Hallmark pathway enrichment (Wk16 vs. Pre) in DPT cells (**N**) or Interferon gamma response pathway enrichment (Wk16 vs. Pre) in classical monocytes (**O**), for each BCG vaccine group. Enrichment score (ES), NES, and BH-adjusted *P* value for IV and ID are given above each plot.

To characterize the transcriptional response to BCG vaccination in PBMCs and BAL at the single cell level, CITE-Seq (scRNA-Seq + surface protein) was performed on samples from Pre and Wk4 and Wk16 post-vaccination (**Figure S2A**). To mimic Mtb exposure, cells were stimulated *in vitro* with WCL before sorting (live, CD45+ leukocytes) for library generation using the 10x Genomics platform. For initial analysis, standard CellRanger and Seurat^31^ single-cell processing pipelines, ambient RNA removal (SoupX^32^) and surface antibody-derived tag (ADT) normalization (DSB^33^) were used. To improve cell type annotation accuracy, an iterative cluster-pool-cluster approach using ADT-facilitated gating was employed, with a focus on T cell and myeloid subsets (**Figures 1B and S2A**). Specifically, ADT-based gating resolved well-documented issues of low CD4 single-cell gene expression in macaques^24,34^ and CD2/T-cell contamination of myeloid populations^35,36^ (**Figures S2B and S2C)**. Clusters were annotated using differentially expressed genes and canonical surface protein expression patterns. Cell types were labeled at multiple clustering resolutions, generated via pooling clusters into broader groups for different types of analyses: *fine* resolution, *intermediate* resolution (gene expression analysis), *broad* resolution (high-level cell frequency analysis) and *all cells* pseudobulk (MDS/PCA analysis) (**Figures 1B and S2A**). Across the 18 macaques and 3 timepoints, 186,952 high-quality single cells in the blood and 201,609 in the BAL were analyzed (**Figure S2D**), comprising a unique reference atlas for ongoing and future work examining rhesus macaque cell-type-specific responses to BCG vaccination.

### Vaccination Phase PBMC Transcriptional Analysis

To evaluate global transcriptional changes in peripheral blood (PBMCs) induced by BCG vaccination, we performed classical multidimensional scaling (MDS, or PCoA), which is mathematically equivalent to PCA analysis in this setting, on pseudobulk counts (*all-cells*) for each sample and timepoint. After correcting for batch and covariate effects, the data separated by BCG vaccination status with samples from IV BCG macaques higher along dimension (Dim)1 than ID BCG macaques at both Wk4 and Wk16 (**Figure 1C**). Dim1 high was characterized by expression of T cell activation genes such as *TNFSF8*, *LCP2*, and *CD247* whereas Dim1 low associated with myeloid-related genes such as *TLR2*, *FCGR2B*, and *IFI30* (**Table S2**). These data reveal IV and ID BCG samples differ in global transcriptional profiles after vaccination, which may be due to cellular composition differences, differential gene expression within cell types, or both.

To further characterize cell-type specific analyses within the PBMC data, Uniform Manifold Approximation and Projection (UMAP) was used to visualize metaclusters of major cell types (**Figure 1D**) and cells were annotated at multiple cluster resolutions (**Figure S3**). We then fit a quasibinomial generalized linear model to determine cell compositional changes after BCG vaccination that may underlie gene expression dynamics. Among all PBMCs, the proportions of T/NK cell, B cell, and monocyte/dendritic cell (mono/DC) metaclusters were generally stable after vaccination other than a transient increase in the T/NK cell metacluster at Wk4 after IV BCG, consistent with prior flow cytometry data^14^ (**Figure 1E**). The T/NK metacluster was partitioned using ADT gating (**Figure 1D**, inset) into a CD4-positive metacluster (CD4+ T cells, including CD4+CD8ɑ+ double-positive T cells; **Figure 1F**) and a CD4-negative metacluster (CD8+ T cells, NK cells, and double-negative T cells; **Figure 1H**). As expected, most T cells in the blood were categorized as naïve or innate-like clusters. After IV BCG, significant increases were observed in polyfunctional (*IFNG, TNF*) CD4 T cells and a population of polyfunctional double-positive T (DPT) cells **(Figures 1G and S3**), as well as increases in polyfunctional CD8 T cells **(Figure 1I**). IV BCG induced minimal compositional changes in the blood mono/DC metacluster compared to Pre except a relative increase in non-classical (*FCGR3*/CD16) monocytes expressing ribosomal genes (e.g., *RPL6*, *RPS3*) from Pre to Wk4 and in conventional dendritic cells (cDCs) from Pre to Wk16 (**Figures 1J and 1K**). Thus, while alterations in the PBMC composition in response to BCG immunization were subtle, IV BCG vaccination generated cytokine-producing polyfunctional T cells earlier and to a greater extent than ID BCG.

To assess variation in cell type-specific gene expression between BCG vaccine groups and timepoints, we used limma^37^ to fit linear models to normalized expression counts, accounting for batch, age, weight, repeated measurements (using duplicateCorrelation), and sample quality heterogeneity. Nine longitudinal contrasts were prespecified for statistical analysis: three timepoint contrasts within each vaccine group (Wk4 vs. Pre; Wk16 vs. Pre; W16 vs. Wk4) and three IV vs. ID differences in longitudinal change. Broadly, the IV BCG dataset comprised a higher proportion of differentially expressed (DE) genes (DEGs) in response to vaccination compared with ID BCG for each timepoint contrast (**Figure 1L and Table S3**). More than half the genes profiled were DE (FDR 0.1) after IV BCG (Wk4 vs. Pre), with many shared among three or more (3+) cell types, suggestive of a coordinated response. Within ID BCG, polyfunctional CD4 T cells displayed the most DEGs among profiled cell types. In comparison, IV BCG elicited DEGs in multiple T cell types including polyfunctional CD4 and CD8 T cells. However, DPT cells displayed the greatest degree of modulation after IV BCG, with 55% of the tested genes DE from Pre to Wk4. When contrasting DEGs from IV vs. ID DPT cells, 28% of genes were differentially responsive from Pre to Wk4 (**Figure 1L**), indicating considerable differences in the effect of each vaccine on DPT cell gene expression.

To uncover the most responsive DPT genes after IV BCG we derived a signature score (“IV-specific DPT signature”), using the 100 most significant DEGs (IV; Wk4 vs. Pre) (**Figure S4A**). Among the 100 genes identified at Wk4, 79 were sustained at Wk16 (**Figure S4B**). Overrepresentation analysis (ORA) using the IV DPT signature gene list revealed enrichment of Hallmark pathways including MYC targets V1, IL2/STAT5 signaling, MTORC1 signaling, and Unfolded protein response, indicating biosynthetic or metabolic activity consistent with activation and proliferation (**Figure S4C)**. DPT cells from IV BCG samples (as well as polyfunctional CD4 and CD8 T cells), had more *Ki67* transcript and protein expressing cells at Wk4 than ID samples consistent with active proliferation (**Figure S4D and Figure S1D**). Compared to ID BCG, DPT cells from IV BCG PBMCs expressed higher (log_2_ FC) *IFNG*, *TNF*, *IL2*, and *RORC* after IV BCG, although these transcripts were not among the 100 *most* significant DEGs (**Figure S4B**). Prior single cell transcriptional analyses identified Th1/Th17 gene modules enriched after IV BCG immunization in BAL (“module 2hi” and “T1-T17”) and in resolving granulomas in lung^11,24,28^. We created modules comprising the top 50 highly expressed genes (log_2_ FC) from these two modules to test within the DEGs of DPT cells profiled here. There was significant enrichment (CAMERA^38^) of modules from both previously defined Th1/Th17 signatures between IV and ID BCG at Wk4 and Wk16 (vs. Pre); however, the IV-specific DPT blood signature defined here captured the greatest differential enrichment between vaccine groups at both timepoints (**Figure S4E**).

To detect coordinated activity of genes across all cell types, gene set enrichment analysis (GSEA) for all 50 Hallmark pathways was performed using the ranked DE gene lists (**Figure S5**). Enrichment of proliferation-related and immune-related pathways induced by ID or IV BCG immunization (vs. Pre) was observed across a broad range of cell types; however, we focused our analysis on the IV vs. ID contrasts over time (**Figure S5**, bottom row). Consistent with the ORA of IV BCG responsive T cells above, DPT cells in IV vs. ID samples at Wk4 and Wk16 showed enrichment of MYC targets V1 and MTOR1 signaling, as well as MYC targets V2, E2F targets, G2M checkpoint, and IL2/STAT5 signaling pathways (**Figures 1M, 1N, and S5**). Together, ORA and GSEA combined with *Ki67* gene and protein expression indicate DPT cells are Th1/Th17 cells with enhanced metabolic activity consistent with an activated and proliferative state.

In assessing the myeloid compartment, GSEA revealed substantial enrichment within blood monocytes for multiple immune pathways, including interferon response pathways (**Figure S5**). Indeed, although few DEGs were detected from APCs after BCG (**Figure 1L**), classical monocytes were enriched in interferon (IFN) alpha (IFNA) and IFNG response pathways to a similar extent from Pre to Wk4 after ID and IV BCG (IV vs. ID; **Figure 1M**). However, from Wk4 to Wk16, IFNA and IFNG signaling continued to increase in IV BCG monocytes, leading to significant enrichment in IV vs. ID by Wk16 (**Figure 1M and 1O**). Thus, both IV and ID BCG vaccination result in transcriptional modulation with enhanced T cell proliferation and IFN signaling in monocytes; however, IV BCG demonstrates a larger and more sustained effect at the later Wk16 timepoint.

### Vaccine Phase BAL Transcriptional Analysis

A similar pipeline (**Figure S2**) was used to analyze BAL samples collected from macaques before (Pre) and after (Wk4 and Wk16) ID or IV BCG vaccination. Gene expression and ADT surface markers for cell type annotation are shown in **Figure S6**. As observed in PBMCs, IV BCG vaccination induced transcriptional changes across pseudobulked BAL data (*all cells*), with IV BCG samples separating from Pre and ID BCG samples (**Figure 2A)**. However, the magnitude of global transcriptional modulation was higher in BAL compared to PBMCs (38.4% vs 18.35% variance in Dim1; **Figures 2A and 1C**), Genes associated with Dim1-high samples included regulators of gene expression, such as *IRF4*, *DAZAP1*, *ILF3*, and *SMARCE1*. In BAL, IV BCG Wk4 and Wk16 samples collected at Wk4 and Wk16 separated along Dim2 (8.9%), suggesting ongoing gene expression modulation throughout the vaccination phase (**Table S2**).

**Figure 2.**
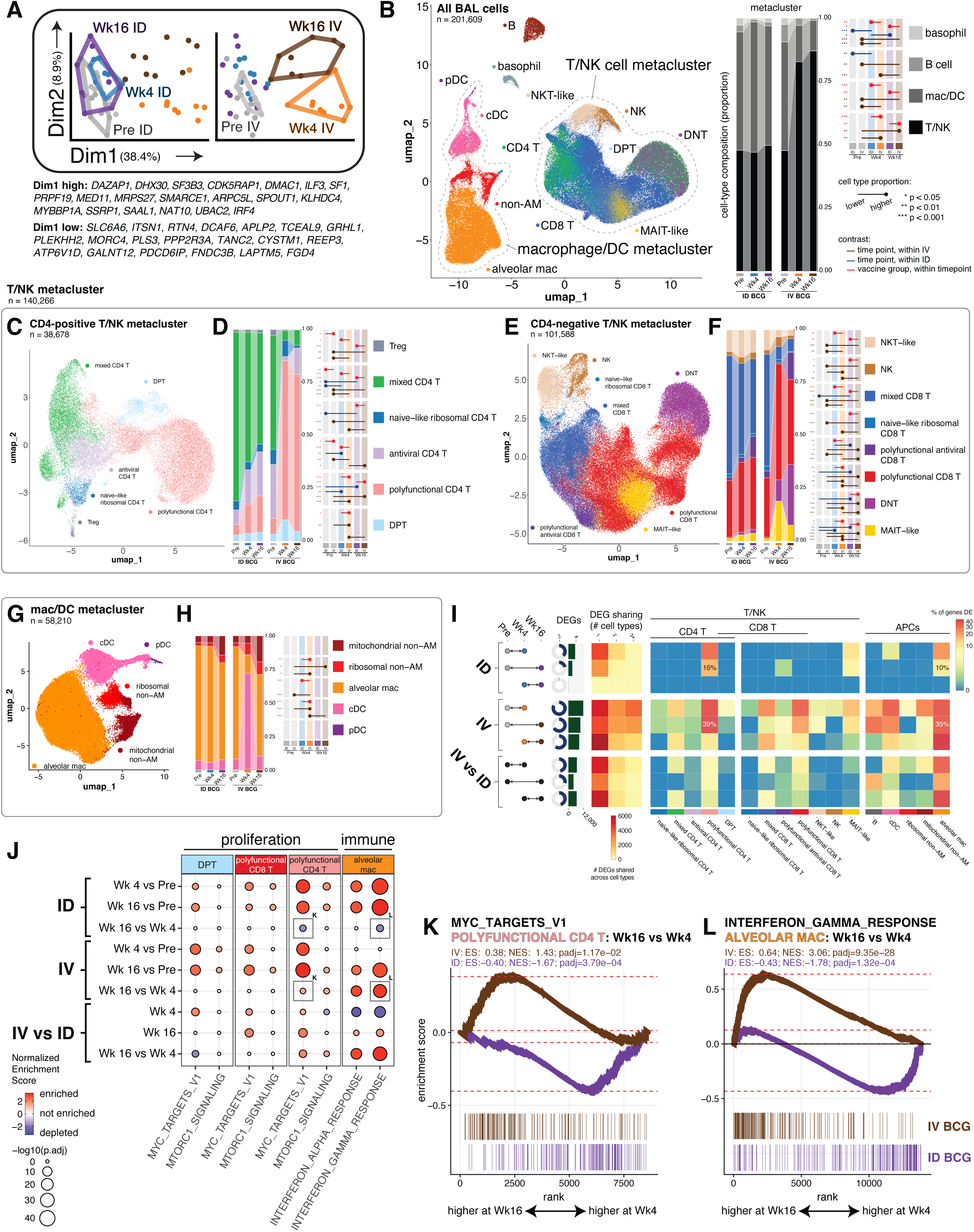
Single-cell CITE-Seq analysis of vaccination phase BAL. **(A)** MDS of BAL data pseudobulked by sample at each timepoint (top). Colored polygons include samples from macaques within a timepoint and vaccine group, ID BCG (left) and IV BCG (right). Top 20 genes associated with Dim1 high and Dim1 low listed below. **(B)** UMAP of scRNA-Seq profiles of vaccination phase BAL (*all cells*) and cell type composition (metacluster proportion of all cells) within ID and IV BCG samples at each timepoint (stacked bars). Significant differences (*P*.adj < 0.05, BH FDR) between timepoints within IV (brown) or ID (blue), or between vaccine groups within timepoint (red) determined by generalized linear modeling are depicted with color-coded directional lollipops (right). (**C**-**H**) UMAP of BAL cells within the CD4-positive T/NK metacluster (**C**), BAL CD4-negative T/NK metacluster (**E**), mac/DC metacluster (**G**), with cell type composition (stacked bars) and significant changes (**D**, **F**, **H**) as in (**B**). **(I)** DE analysis of genes in BAL for each timepoint contrast within ID (top) or IV (middle), or between vaccine groups across timepoints (bottom). Percent (donuts) and number (bars) of DEGs (of all genes profiled) across all cell types and DEG sharing heatmap depicting the number of DEGs (color scale below) shared across 1, 2, or 3 (or more) cell types. Within the T/NK cells and APCs (B cells, mac/DCs), heatmap illustrating the percent of DEGs (color scale on right) within each cell type (labeled below) for each contrast. Significant differential expression was defined as FDR < 0.1 (BH). **(J)** Selected GSEA results. NES and associated *P* values based on GSEA of rank-ordered signed *P* value lists (−log10*P* × sign(logFC)) derived from DE analysis. Shown are selected enriched Hallmark pathways (below) within polyfunctional CD4 T, polyfunctional CD8 T, DPT cells, and alveolar macrophages for each contrast (left). Complete GSEA is presented in **Figure S8**. (**K**-**L**) Barcode plots of MYC targets v1 pathway enrichment (Wk16 vs. Wk4) in polyfunctional CD4 T cells (**K**) or Interferon gamma response pathway enrichment (Wk16 vs. Wk4) in alveolar macrophages (**L**), for each BCG vaccine group. Enrichment score (ES), NES, and BH-adjusted *P* value for IV and ID are given above each barcode plot.

IV BCG induced substantial changes in BAL cell composition compared to ID BCG (**Figure 2B**). Consistent with flow cytometry analysis (**Figure S1A**), the proportion of cells in the T/NK cell metacluster increased from Pre to Wk4 after IV BCG vaccination, whereas the relative proportion of macrophage/DCs decreased (**Figure 2B**). Notably, within the CD4-positive (**Figure 2C and 2D**) and CD4-negative metacluster (**Figure 2E and 2F**), timepoint-associated and vaccine group-associated effects were observed in every annotated cell type. Polyfunctional CD4 and CD8 T cells increased in BAL after ID and IV BCG; however, proportions of these cells were ∼5 fold (CD4) and ∼1.5 fold (CD8) higher in IV compared to ID BCG samples at Wk4 and Wk16 (**Figure 2D**). The proportion of DPT cells in IV BCG BAL samples increased to a lesser extent than polyfunctional CD4 T cells but was significantly higher than Pre and ID BCG at Wk4. Compared with ID, IV BCG elicited more double-negative T (likely gamma delta T cells that did not cluster separately here by gene expression or ADT) and MAIT-like T cells (both of which showed *IFNG* and *TNF* expression, with or without *IL2*, **Figure S6**) at Wk4 (**Figure 2F**). Within the substantially reduced IV BCG macrophage/DC metacluster (**Figure 2G and 2H**), there was a transient significant increase in the proportion of conventional dendritic cells (cDCs) compared to AMs (Wk4 vs. Pre). There was a small but significant increase in ribosomal non-AM (*CXCL10*/*11*+ *MARCO*-), which may be analogous to recruited macrophages, reported to be increased relative to AMs in the BAL after IV BCG in prior work^28^. By Wk16, proportions of both myeloid subsets returned to near-baseline levels (**Figure 2H**).

DE gene analysis via linear modeling was then performed on the BAL dataset (**Figure 2I** and **Table S3**). IV BCG induced broader and more dynamic DE in the BAL than ID BCG, with vaccine-responsiveness observed in nearly all cell types and timepoints. Up to 80% of all genes tested were DE (FDR 0.1) Wk4 after IV BCG compared to 30% for ID (**Figure 2I**, donut plots). Cell-type specific analysis showed polyfunctional CD4 T cells and AMs were highly responsive to ID and IV BCG although IV-induced changes from baseline were of greater magnitude and more persistent. From Pre to Wk16, 39% of genes were DE in polyfunctional CD4 T cells after IV BCG (compared to 16% after ID BCG) and 35% of genes were DE in AMs (10% after ID) (**Figure 2I**). Polyfunctional CD8 T and DPT cells were also responsive to IV BCG but to a lesser extent than polyfunctional CD4 T cells. Among APCs, B cells, cDCs and non-AM had significant DE, but AMs substantially increased in DEGs from Wk4 to Wk16. (**Figure 2I**). Of note, among all cell types, DE analysis revealed that gene expression in AMs was by far most divergent between IV and ID, especially for the Wk16 vs. Wk4 contrast. Among the DEGs in AMs, the top overrepresented Hallmark pathways included the IFNG response, IFNA response, and IL6/JAK/STAT3 signaling pathways (**Figure S7).**

GSEA for Hallmark pathways was next performed to characterize coordinated expression of pathway genes across all cell types in BAL (**Figure S8 and Table S3**). Generally, there was enrichment of various immune, metabolic, and proliferation-related pathways in many cell types after both vaccines for each contrast. As it was the timepoint closest to Mtb challenge, we focused further BAL analysis on cell types with pathway enrichments at Wk16. Polyfunctional CD4, CD8 and DPT cells were enriched for pathways associated with proliferation (MYC Targets and/or MTORC1 Signaling) at Wk4 (vs. Pre) after IV and ID BCG vaccination (**Figure 2J**). However, there was continued enrichment of proliferation pathways in IV samples from Wk4 to Wk16, while these pathways decreased in ID samples (**Figures 2J and 2K**). Within AMs, genes associated with multiple immune pathways (including IFNA and IFNG response pathways) were enriched at Wk4 in ID vs. IV samples (**Figures 2J and S8)**. However, between Wk4 and Wk16, IFNA and IFNG response pathway genes were upregulated in AMs from IV samples, leading to greater pathway enrichment in IV vs. ID samples at Wk16, consistent with ORA results (**Figures 2J**, **2L, and S7**). Together, these data suggest IV BCG specifically promotes sustained upregulation of proliferation pathways in polyfunctional cytokine-producing T cells and IFN response pathways in BAL AMs from Wk4 to Wk16 compared to ID BCG.

### Analysis of Inferred Cellular Communication Networks in BAL

A key finding from the BAL transcriptional analysis is the sustained IFNG response in AMs following IV BCG that may be functionally important for TB control. Indeed, following IV BCG, IFNG production from CD4 T cells after IV BCG is critical to maintain IFNG responsiveness and control of lung viral infections in mouse models^19–21^. To characterize the effect of BCG immunization on cell-cell communication networks in BAL, we used CellChat^39^ to map sender and receiver cells for ligand-receptor interactions across 17 cell types, focusing deeper on T cell interactions with AMs. Across CellChat pathways and cell types, IV BCG induced more inferred pathway interactions and greater aggregate pathway strength than ID BCG at both timepoints (**Figure 3A**). ID BCG increased interaction counts relative to Pre, but IV BCG increased both counts and strength. This indicates broadly enhanced cell-cell communication with IV vs. ID BCG, likely underlying the transcriptional changes observed in BAL DE and Hallmark GSEA analyses (**Figure 2I)**.

**Figure 3.**
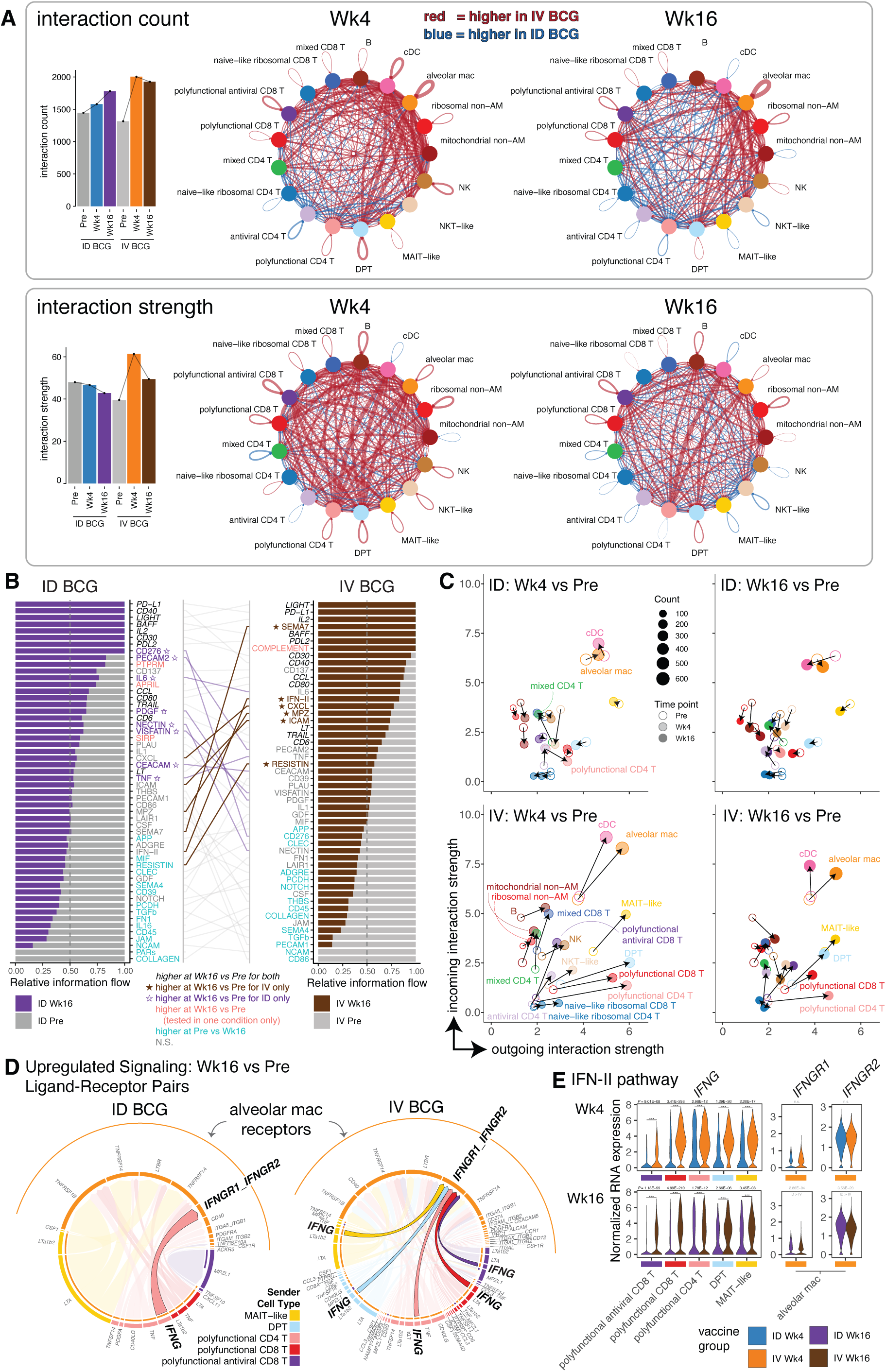
Cell-cell interaction analysis of vaccine-phase BAL. **(A)** Ligand-receptor interaction count (top) and interaction strength (bottom) across CellChat pathways and cell types (CellChat permutation test for interaction *P* < 0.05) at each timepoint for IV and ID BCG BAL samples (bars, left). Differences in inferred cell type-to-cell type interaction count and strength across all pathways at Wk4 and Wk16 post-vaccination (network plots, right). Line thickness reflects the relative magnitude of the difference between IV (red) and ID (blue) BCG in count or strength. **(B)** Relative information flow of CellChat pathways at Wk16 vs. Pre (Wk4 vs. Pre; **Figure S9A**), for ID and IV BCG across all cell types in BAL. Center lines connect matching pathways; stars indicate those significantly induced in either ID only (purple) or IV only (brown). \**P* < 0.05, CellChat rankNet test. **(C)** Vector plots of incoming and outgoing signal strength (summed for all pathways), for each cell type at Wk4 vs. Pre (left) and Wk16 vs. Pre (right) in ID BCG (top) and IV BCG (bottom). Arrows connect same cell type from Pre-to post-BCG. **(D)** CellChat-inferred IFN-II (IFNG) pathway communication between T cells and AMs in BAL from ID or IV BCG samples upregulated at Wk16 vs. Pre. Chords connect ligand gene-expressing source cells with receptor gene-expressing target cells and require significant ligand upregulation (Wilcoxon rank-sum test, *P* <0.05; logFC > 0.05). Full CellChat pathway analysis (Wk16 vs. Pre) between T cells and AMs is shown in **Figure S9C**. **(E)** Violin plots of normalized RNA expression for IFN-II pathway ligand (*IFNG*) and receptors (*IFNGR1*/*2*) in T cell subsets and AMs from ID and IV BCG samples at Wk4 and Wk16. *P* values indicate IV vs. ID differences (Wilcoxon rank-sum test); n.s., not significant.

We next performed a relative information flow analysis to identify pathways, across all cell types, significantly induced after IV vs. ID BCG at Wk4 (**Figure S9A**) and Wk16 (**Figure 3B**). From Pre to Wk16, many pathways showed increased signaling in both groups, including immune-modulating pathways CD40, LIGHT, PD-L1, and IL2 (**Figure 3B**). CellChat pathways significantly upregulated in IV but not ID at Wk16 included SEMA7, which can promote myeloid cytokine production via cell-cell contact^40^, as well as IFN-II (IFNG) and CXCL chemokine pathways. Pathways induced in ID but not IV included CD276, which can constrain T cell proliferation^41,42^, and inflammation-related pathways IL6 and TNF. Beyond simple quantitative differences in interaction count and strength, IV and ID showed substantial differences in their predominant communication networks. Of note, CellChat did not detect IFN-I (IFNA) pathway activity in this dataset due to insufficient direct ligand evidence. IFNA transcripts are intrinsically sparse and difficult to quantify reliably from the highly homologous multigene IFNA locus using short-read scRNA-seq; however, this does not preclude analysis of downstream IFNA signaling in target cells. (see methods).

To identify the major cell types in the BAL driving global changes in pathway signaling after ID or IV BCG, we visualized inferred incoming and outgoing interaction strength for each cell type from Pre to Wk4 and Wk16 (**Figure 3C**). IV BCG macaques showed increased signaling strength across a diversity of cell types at Wk4. At Wk16, there were sustained increases in outgoing signaling strength from MAIT-like, polyfunctional CD4, CD8, and DP T cells, as well as sustained incoming signal strength to AMs and cDCs. In ID animals, all cell types remained close to their baseline at both timepoints, with only the polyfunctional CD4 T cell population transiently increasing outgoing signaling strength at Wk4 (**Figure 3C**). Thus, while both BCG routes alter signaling network quantity and strength across cell types, IV BCG elicits greater and more persistent shifts, with notable sustained patterns in polyfunctional T cells, cDCs, and AMs.

Given AM gene expression and communication patterns differed extensively between IV and ID vaccinated animals, we next focused on sustained changes in the ligand-receptor expression and communication probability between T cells and AMs in BAL (**Figure S9B**). Prior analyses reported increased aggregate pathway strength of T cell to AM interactions in BAL from IV vs. ID BCG but did not evaluate individual pathways, cell types, or timepoints for BCG route-specific differences^28^. At the ligand-receptor level, IV BCG vaccination elicited broader and qualitatively distinct remodeling of bidirectional AM–T cell communication in BAL at Wk16 (vs. Pre) compared to ID BCG. Significant communication probability from AMs to T cells in IV samples was characterized by pathways of cell adhesion, co-stimulation, and proinflammation (ICAM, CD80, RESISTIN) whereas ID vaccination was more prominently associated with increased AM-derived TNF signaling (**Figure S9B and S9C**). Inferred communication from T cells to AMs after IV BCG vaccination was broad and included pathways of activation, co-stimulation, and recruitment (IFN-II, LIGHT, CD40, and CCL). Analysis of interactions associated with ligand expression showed that T cell signaling to AMs in ID (Wk16 vs. Pre) was largely dominated by MAIT-like cells sending LTA to AMs, whereas IV displayed a greater variety of sender cells with overlapping signaling activity (**Figure S9C**). A striking example was IFN-II signaling from T cells to AMs; in ID samples only polyfunctional CD4 T cells sent IFNG to AMs, whereas in IV samples, polyfunctional (DPT, CD4, and CD8), antiviral polyfunctional CD8, and MAIT-like T cells sent IFNG to AMs. (**Figure 3D**).

Increased pathway activity can result from increased ligand expression on sender cells and/or increased receptor expression on receiver cells. At Wk4 and Wk16, *IFNGR1* and *IFNGR2* gene expression was similar between AMs from IV and ID animals, while *IFNG* gene expression across selected T cells was higher in IV than in ID at both timepoints (**Figure 3E**). Consistent with this transcriptional data, IFNG protein levels from WCL-stimulated BAL and PBMC cell supernatants were higher in IV vs. ID at Wk4 and Wk16 post-BCG (**Figure S10**). Other pathways with higher inferred activity in IV AMs, such as LIGHT and CD80, showed higher receptor gene expression in IV vs. ID AMs as well as increased ligand expression in source cells (**Figure S9D).** These data suggest the sustained IFNG response pathway enrichment (ORA and GSEA, **Figures S7 and 2L**) in BAL AMs from IV vs. ID immunized macaques may be mediated by a greater variety of T cell types expressing higher IFNG after IV BCG.

### Challenge Phase Outcome and CITE-Seq Analysis

Until now, there has been limited transcriptional analysis of the initial vaccine-induced immune responses after Mtb infection–when AMs in BAL are responding to infection and migrating into lung, and when adaptive responses are being recalled. Across several prior studies, >50% of rhesus macaques immunized with IV BCG showed no evidence of infection in lungs 8-12 weeks after Mtb challenge and no detectable primary T cell responses to Mtb-specific antigens (ESAT-6, CFP10) suggesting very early control of TB^11,12,14^. In contrast, ID BCG at the clinically approved human dose used here, or at higher doses matching IV BCG, confers substantially less protection in macaques than IV BCG^11^. These findings suggest that IV (but not ID) BCG induces an immune response capable of rapidly eliminating Mtb and preventing disease. *In vivo* depletion studies showed that IV BCG-mediated control of TB disease is dependent on CD4 and CD8ɑ expressing cells but may also involve enhanced function of AMs since they are the first target of Mtb infection^22,43–45^. This hypothesis is supported by data from prior efficacy studies in which macaques were challenged with a barcoded strain of Mtb Erdman that allowed infection establishment to be tracked; IV BCG restricted the number of individual Mtb bacilli (barcodes) that establish infection compared to unvaccinated animals, and this early restriction was largely maintained in T cell-depleted, IV BCG macaques even though endpoint Mtb burdens were high (**Figure S11**). Unlike after IV BCG, no early reduction of Mtb bacilli (barcodes) was observed after challenge of ID BCG animals (**Figure S11A**). We hypothesized that transcriptional patterns in the airway and lung in response to Mtb challenge may define distinct pathways underlying early Mtb restriction by activated AMs after IV, but not ID BCG immunization.

To characterize the initial transcriptional response post-challenge at the site of infection (airway and lung), a subset of ID and IV BCG immunized macaques (n=3 per vaccine group and 3 unvaccinated controls) were infected 22 weeks after BCG (**Figure 4A** and **Table S1**). For this study, we used fluorescently labelled Mtb Erdman mCherry (16–37 CFU) to collect imaging data for a separate analysis, rather than barcoded Mtb Erdman. To evaluate early control of infection, Mtb burdens (CFU) were measured in lung tissue and lung lymph nodes D11 post-challenge (**Figure 4B**). Mtb burdens in lung and lung lymph nodes trended lower after IV BCG compared macaques to ID BCG and unvaccinated controls, although statistical significance cannot be achieved with 3 animals per group.

**Figure 4.**
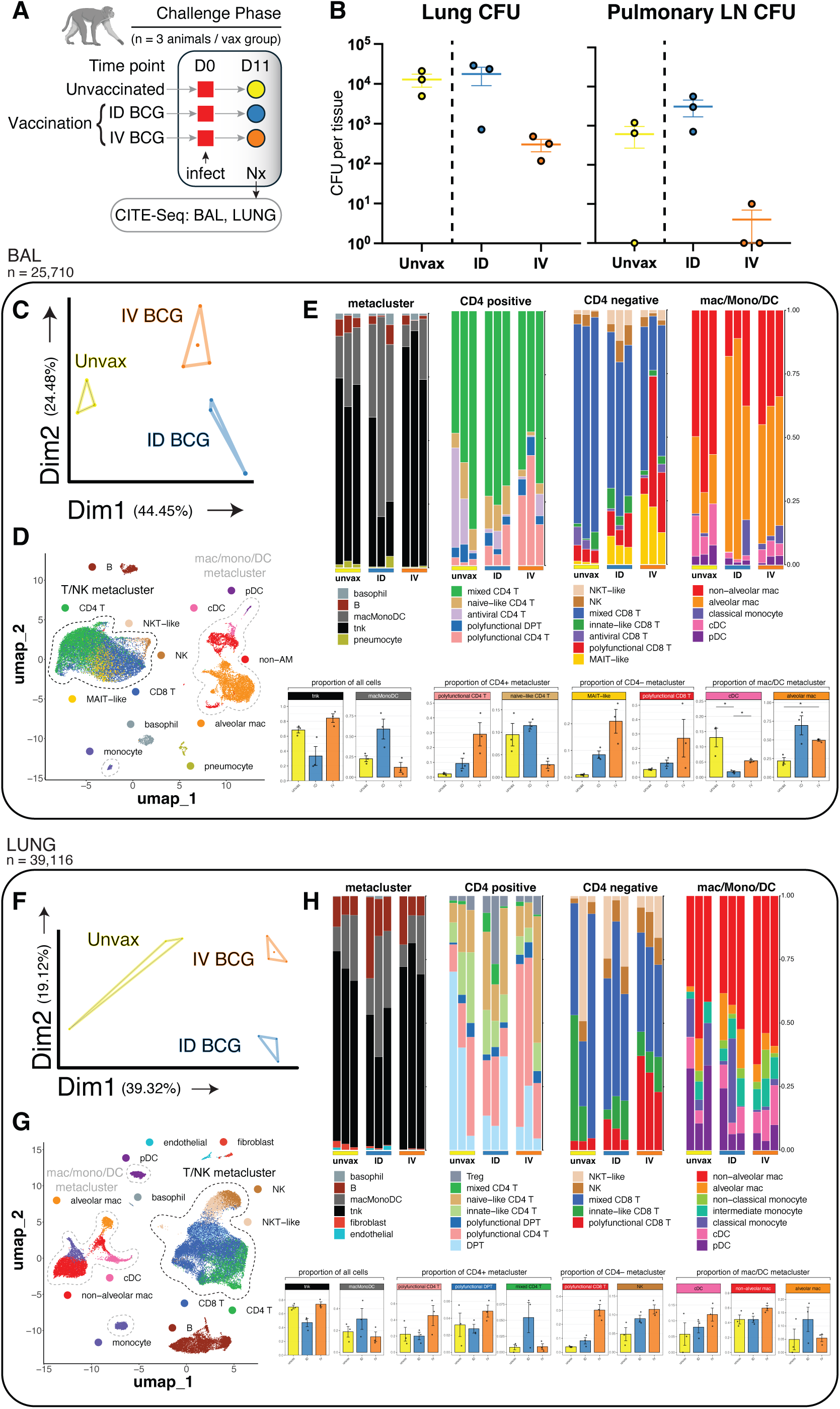
Mtb challenge phase and CITE-Seq analysis of BAL and lung after infection. **(A)** Three macaques per group (unvaccinated, ID BCG, IV BCG) were infected with Mtb Erdman (16-37 CFU) 22 weeks after vaccination (D0); BAL and lung tissue were collected and processed 11 days (D11) after infection. **(B)** Mtb bacterial burdens D11 after infection in the lung and pulmonary lymph nodes presented as total CFU per total amount tissue processed. (**C**, **F**) MDS of post-challenge BAL (**C**) and lung cells (**F**) pseudobulked data (*all cells*). Colored triangles include samples from macaques within a vaccine group; IV BCG group includes one technical replicate across 10x processing batches. (**D**, **G**) UMAP of scRNA-Seq profiles of BAL (**D**) and lung (**G**) D11 after infection. (**E**, **H**) Cell type composition (proportion of cell types) in BAL (**E**) and lung (**H**) for *all cells* and for the CD4-positive T/NK metacluster, CD4-negative T/NK metacluster, and mac/mono/DC metacluster. Data from individual animals are presented as stacked bars (cell types are color-coded below), and as bar charts (color coded by vaccine group) for statistical testing (*P*.adj < 0.05, BH FDR).

CITE-Seq data was generated from cryopreserved BAL and randomly selected lung tissue without additional *in vitro* stimulation since these samples were collected from *in vivo* infected tissues **(Figure S12A)**. The post-challenge analysis comprised fewer animals per condition and fewer profiled cells per animal and after QC, we analyzed libraries of 25,710 BAL and 39,023 lung cells (**Figure S12B**). Clustering and cell type annotation for the challenge phase data was performed similar to but separate from, the vaccination phase data (**Figure S13**).

MDS analysis of post-challenge data (*all cells*) showed substantial global transcriptional changes between the three groups of macaques in BAL and lung (**Figures 4C and 4F, Table S2**). In both tissues at D11 post-infection, Dim1 separated unvaccinated from vaccinated samples while Dim2 separated IV vs. ID BCG samples. Analysis of cell proportions within the BAL (**Figures 4D and 4E**) and lung (**Figures 4G and 4H**) showed differences between IV and ID samples similar to the pre-challenge phase BAL samples. Specifically, the ratio of T/NK cells to myeloid cells (mac/mono/DC) was higher in IV BCG animals compared to ID BCG. Of note, in unvaccinated macaques, T/NK proportions more closely reflected IV than ID vaccinated animals, although IV animals had the highest proportions of polyfunctional T cells in both compartments (**Figures 4E and 4H**). Overall, these data suggest that vaccine-induced changes in cell composition between IV and ID in BAL prior to challenge were recapitulated after Mtb infection in BAL and lung.

DE analysis focused on IV vs. ID contrasts in BAL and lung for T cells and APCs (**Figure S14A, Table S3**). Overall, there was a lower percentage of significant DEGs in the post-challenge compared to the pre-challenge dataset, potentially due to less antigen stimulation from natural TB infection *in vivo* compared to *in vitro* stimulation, or due to lower statistical power to detect differences between smaller sample numbers. Therefore, we proceeded directly to GSEA to detect coordinated pathways enriched among ranked gene lists from the DE analysis (**Figures 5A and S14B**). At D11 post-challenge in BAL, immune-related pathways, including IFNA and IFNG response pathways, inflammatory response and TNF signaling were enriched across nearly all APCs in IV vs. ID BCG vaccinated animals. This was consistent with pre-challenge BAL data, which showed substantial enrichment of the IFNA and IFNG response pathways from Wk4 to Wk16 after vaccination in IV vs. ID BAL AMs (**Figure 2J**). Lung tissue differed from the BAL in post-challenge samples with most immune-related pathway enrichment observed in APCs from ID BCG animals compared to IV (**Figure 5A**). This could be due to increased Mtb burden in the lung tissue from ID vs. IV macaques, as monocyte-derived cells from lung granulomas with higher Mtb burdens are skewed toward a more inflammatory response^27^. The exception to this was the enrichment of IFNA and IFNG response pathways in AMs from IV animals, concordant with pre-challenge and post-challenge BAL samples (**Figures 5A and 5B**). These data indicate that while immune pathway enrichment varies between BAL and lung tissue, a consistent finding across both compartments is the enrichment of IFN response genes in AMs from IV BCG animals after challenge.

**Figure 5.**
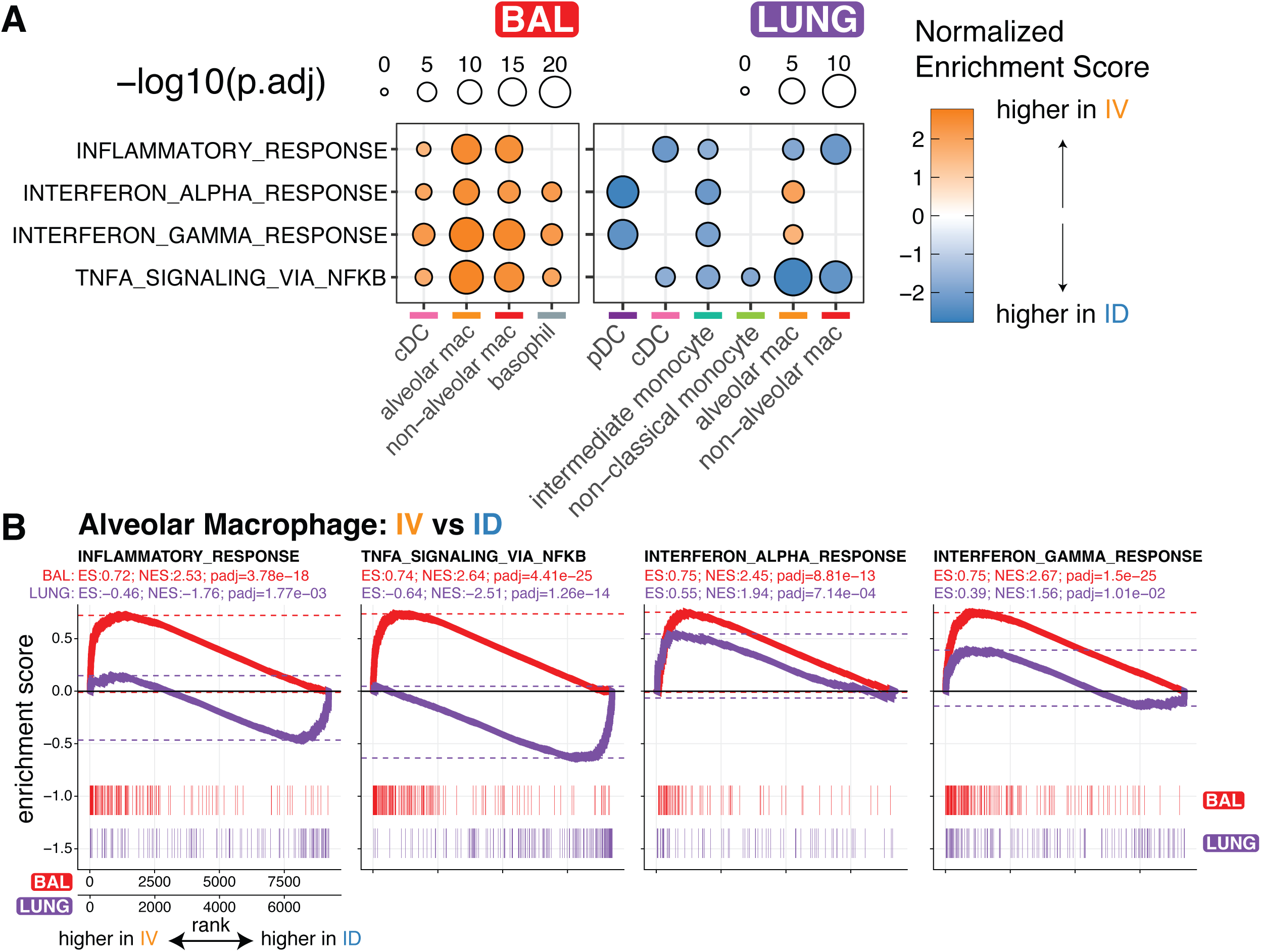
GSEA of BAL and lung cells after Mtb infection. (**A**, **B**) NES and associated BH-adjusted *P* values based on GSEA of rank-ordered signed *P* value lists (−log10*P* × sign(logFC)) derived from DE analysis in **Figure S14A**. Enrichment (IV vs. ID) of selected Hallmark pathways in mac/mono/DC cell types are presented for BAL and lung (**A**). Complete post-challenge GSEA is presented in **Figure S14B**. (**B**) Pathway enrichment (IV vs. ID) of the Inflammatory response, TNFA signaling via NFKB, IFNA response, and IFNG response after challenge in AMs from BAL (red) and lung (purple). Enrichment score (ES), NES, and BH-adjusted *P* value for each tissue are given above each plot.

### Defining a transcriptional correlate of IV BCG-mediated protection against TB

Enrichments of IFNA response and IFNG response pathways were observed in AMs from IV BCG relative to ID BCG during the early response to Mtb challenge (**Figure 5B**). These enrichment patterns emerged prior to challenge, driven by strong increases in IFN response pathway genes in AMs between Wk4 and Wk16 after IV BCG vaccination and a decrease in IFN responses in AMs from ID BCG (**Figure 2L**). Given that most genes in the Hallmark IFNA response pathway (66 of 86 profiled) are shared with the IFNG response pathway (179 genes), we assessed whether the IFNA response signal at Wk16 vs. Wk4 was driven by genes shared between both pathways or was unique to IFNA response genes (**Figure 6A**). This analysis found the shared IFNA and IFNG response genes accounted for more of the IFNA response pathway signal compared to IFNA response unique genes, suggesting IFNG production may contribute to the enriched IFNA response signal pre-challenge. The role of *IFNG* transcript is supported by protein data showing high IFNG production (and low IFNA) between IV and ID BAL and PBMCs at Wk4 and Wk16 post-BCG, in response to WCL stimulation (**Figure S10**). However, because multigene IFNA transcripts are not reliably mapped in rhesus transcriptional datasets and IFNA protein is difficult to measure, a role for Type I IFN cannot be ruled out.

**Figure 6.**
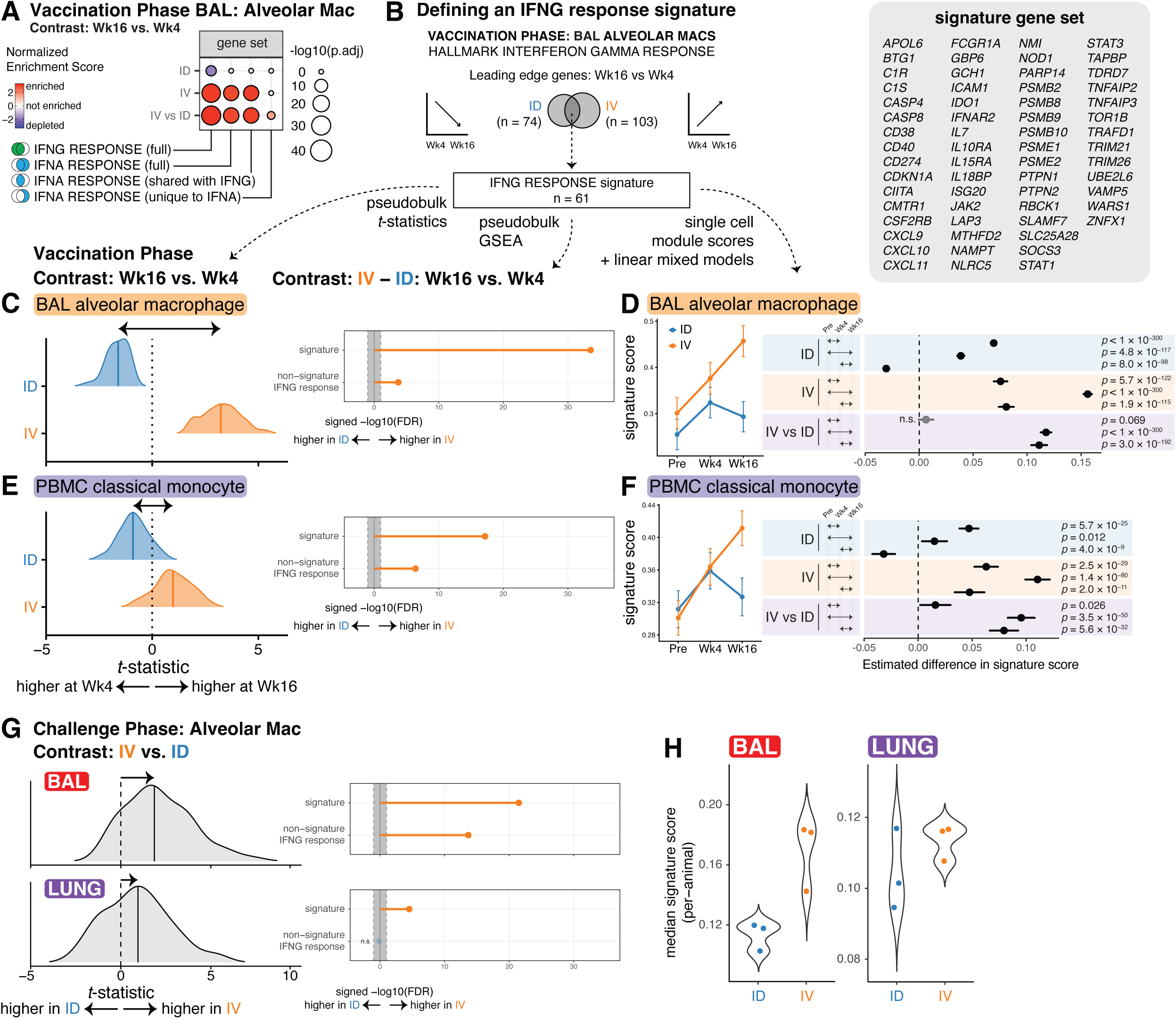
Defining an IFNG response pathway signature in vaccination-phase Ams. **(A)** GSEA (NES and BH-adjusted *P* values) using BAL AM ranked gene lists (Wk16 vs. Wk4: within vaccine groups and between IV and ID) showing enrichment in the IFNG or IFNA response Hallmark (full) gene sets, and enrichment in subsets of genes shared between both IFN response pathways or unique to the IFNA response. **(B)** Schematic for defining and testing a 61-gene IFNG response signature whose expression diverges in AMs from IV vs. ID BCG AMs between Wk4 and Wk16 after BCG immunization; gene set is listed in grey box. (**C**, **E**) Ridgeline density plots (left) showing *t* statistics (direction and strength of change in gene expression) from pseudobulk DE analysis of the 61-gene signature in BAL AMs (**C**) and PBMC classical monocytes (**E**) from ID and IV immunized macaques between Wk4 and Wk16 after BCG. Positive *t* statistics indicate higher expression at Wk16, and negative values indicate higher expression at Wk4; arrows depict difference of *t* statistic medians (solid lines) between vaccine groups. Right, statistical analysis (CAMERA competitive gene tests, lollipops) of enrichment (IV vs. ID) of the IFNG signature response and IFNG non-signature response from Wk4 to Wk16; dotted lines represent significance threshold (BH FDR 0.1 (|signed −log10(FDR)| = 1). (**D**, **F**) Single-cell module scores (UCell) for the 61-gene signature in BAL AMs (**D**) and PBMC classical monocytes (**F**). Left, line graphs show model-adjusted estimated marginal means (EMMs) and 95% confidence intervals at Pre, Wk4, and Wk16 resulting from linear mixed-effects models. Right, forest plots for model-estimated contrasts; dots are estimates of differences based on each contrast and bars indicate 95% confidence intervals. BH-adjusted *P* values (*P.*adj) were calculated across nine pre-specified contrasts, evaluated separately within each tissue. n.s., not significant. **(G)** Direction and strength of difference in IFNG response signature gene expression (gene level *t* statistics) from pseudobulk DE analysis between AMs from IV vs. ID BCG vaccinated macaques D11 after *Mtb* challenge (BAL, top; lung, bottom). CAMERA competitive gene set tests (lollipops) show enrichment (IV vs. ID) of the signature and non-signature IFNG response in post-challenge BAL and lung AMs with significance at FDR 0.1 (dotted lines). **(H)** Violin plot showing the distribution of the median single-cell signature scores for AMs in BAL and lung in both vaccine groups at D11 post *Mtb* challenge. Points represent individual animals.

As IV BAL AMs showed enrichment of the IFNG response pathway at Wk16 vs. Wk4 while ID AMs showed depletion (negative enrichment) at Wk16 vs. Wk4 after vaccination (**Figure 2L**), we assessed the leading-edge genes (LEGs) that drove the diverging enrichments between IV and ID BCG, with the goal of generating a candidate transcriptional signature for predicting TB protection. We found 61 LEGs from the IFNG response pathway strongly contributed to both the positive enrichment of the IFNG response in IV AMs and the negative enrichment of the same response in ID AMs from Wk4 to Wk16, thus defining this gene set as a myeloid IFNG response signature with temporally diverging expression between IV and ID (**Figure 6B**). The 61-gene signature includes genes induced by IFNG receptor binding (*JAK2, STAT1*) as well as genes associated with antigen presentation (*CIITA, TAPBP, NLRC5*), chemokines for T cell and NK cell recruitment (*CXCL9, CXCL10, CXCL11*), and genes associated with T cell co-stimulation and adhesion (*CD40, ICAM1*). Of note, the signatures also included genes associated with inflammatory regulation, such as the metabolic modulators *IDO1* and *GCH1* as well as the checkpoint inhibitor *CD274*. We evaluated the IFNG response signature in BAL at the pseudobulk level and found significant divergent enrichment in IV vs. ID AMs from Wk4 to Wk16 that was greater than the non-signature IFNG response genes (**Figure 6C and Figure S14C**). To complement the pseudobulk analysis, and to directly evaluate the signature across timepoints and vaccine groups in individual AMs, we calculated single-cell module scores (UCell) for the signature gene set and used linear mixed-effects models with covariates (age, weight, batch/10X run block) and a random intercept for animal ID to account for repeated measures. We determined the estimated difference in signature score and *P* value for nine prespecified longitudinal contrasts used throughout (**Figure 6D**). From Pre to Wk4 after BCG, the signature score in both IV and ID AMs increased (IV, *P*adj = 5.7 × 10^−122^; ID, *P*adj < 1 × 10^−300^) and were not significantly different from each other (IV vs. ID, *P*adj = 0.069), indicating that the signature score increased to the same extent in both vaccine groups. However, from Wk4 to Wk16 the 61-gene signature score continued to increase in IV AMs (*P*adj = 1.9 × 10^−115^), while sharply retracting in ID AMs (*P*adj = 8.0 × 10^−98^).

In addition to defining a protective correlate in BAL (the site of infection), extending this correlate to blood would inform clinical vaccine development, as blood is the standard site for immune analysis following vaccination. As our dataset included matched timepoints from BAL and PBMCs, we assessed the IFNG response signature in monocytes from PBMCs during the BCG vaccination phase. As observed in the BAL, at the pseudobulk level, expression of the 61-gene IFNG response signature diverged (IV vs. ID) in classical and non-classical monocytes from Wk4 to Wk16 after BCG and displayed more significant enrichments compared with the non-signature IFNG response genes (**Figures 6E and S14C**). At the single-cell level, we also found that the IFNG response signature score substantially increased in IV classical monocytes from Wk4 to Wk16 (*P*adj = 2.0 × 10^−11^) while decreasing in ID classical monocytes between the same timepoints (*P*adj = 4.0 × 10^−9^) (**Figure 6F)**.

To determine whether the IFNG response signature defined in the post-vaccination dataset was also enriched in IV BCG macaques early after Mtb challenge at the site of infection, we compared AMs from IV vs. ID BCG macaques in the post-challenge BAL and lung data (**Figures 6G and 6H**). The 61-gene signature was significantly enriched (IV vs. ID; pseudobulk) in BAL AMs as well as in lung AMs, albeit with weaker statistical evidence (**Figure 6G**, right); the median signature scores per animal (single cell) were descriptively higher (IV vs. ID) for both BAL and lung (**Figure 6H**). Together, these data show the 61-gene IFNG response signature defined here in BAL AMs is detectable in the peripheral monocytes post-vaccination, reflecting a sustained activated myeloid transcriptional program that may influence protective immunity against TB in the airway after challenge.

### Independent cohort for correlating 61-gene IFNG response signature with protection

The primary “discovery” dataset analyzed in the prior figures defined an IFNG response signature whose expression in myeloid cells temporally diverged in macaques vaccinated with IV vs. ID BCG. To investigate whether the 61-gene signature would associate with protection irrespective of vaccine dose and route, a secondary orthogonal dataset was generated from an independent cohort of IV BCG immunized macaques that underwent an 8-week Mtb challenge phase.

CITE-Seq data was generated from samples collected during a previously published challenge study in which rhesus macaques (n=12) were immunized with IV BCG (∼5 x 10^7^ CFU) for 6 months, challenged with Mtb, and evaluated for protection 8 weeks later^14^. This dataset provides an opportunity to assess the association of transcriptional signatures in IV BCG-immunized macaques with protection through a full course of infection. In this analysis we also assessed whether *in vitro* antigen stimulation influenced the defined signature. Thus, data was generated from both unstimulated and WCL-stimulated PBMC collected before (Pre) and after (Wk4, Wk13, Wk20) IV BCG immunization (**Figure 7A**). To ensure annotation consistency, PBMC data from the independent cohort was reference-mapped onto the UMAP of the PBMCs from the discovery cohort based on transcriptional profiles, and ADT protein expression was used to annotate CD4, CD8, DN, and DP T cell clusters from 530,687 cells (**Figures 7B and 7C**). Cellular proportions were not substantially affected by antigen stimulation, although this could be influenced by reference-mapping to the discovery cohort PBMCs, which were all stimulated. The only difference was a small shift toward a larger proportion of non-classical monocytes after stimulation (**Figure 7C**). Broadly, changes in CD4-positive and CD4-negative metaclusters after IV BCG immunization observed in the discovery cohort were reproduced in the independent cohort. This included increased proportions of polyfunctional CD4 and CD8 T cells at Wk4 after vaccination as well as a sustained increase in the proportion of DPT cells through Wk20. As expected, WCL-stimulation enhanced cytokine (*IFNG, IL2, TNF, IL-17A*) and effector molecule (*GZMB, PRF1*) transcription in these expanded T cell subsets (**Figure S15A**). Overall, these results indicate that the orthogonal dataset was representative of the discovery dataset with respect to cellular composition and that stimulation enhanced sensitivity for detecting antigen-specific T cells in PBMCs.

**Figure 7.**
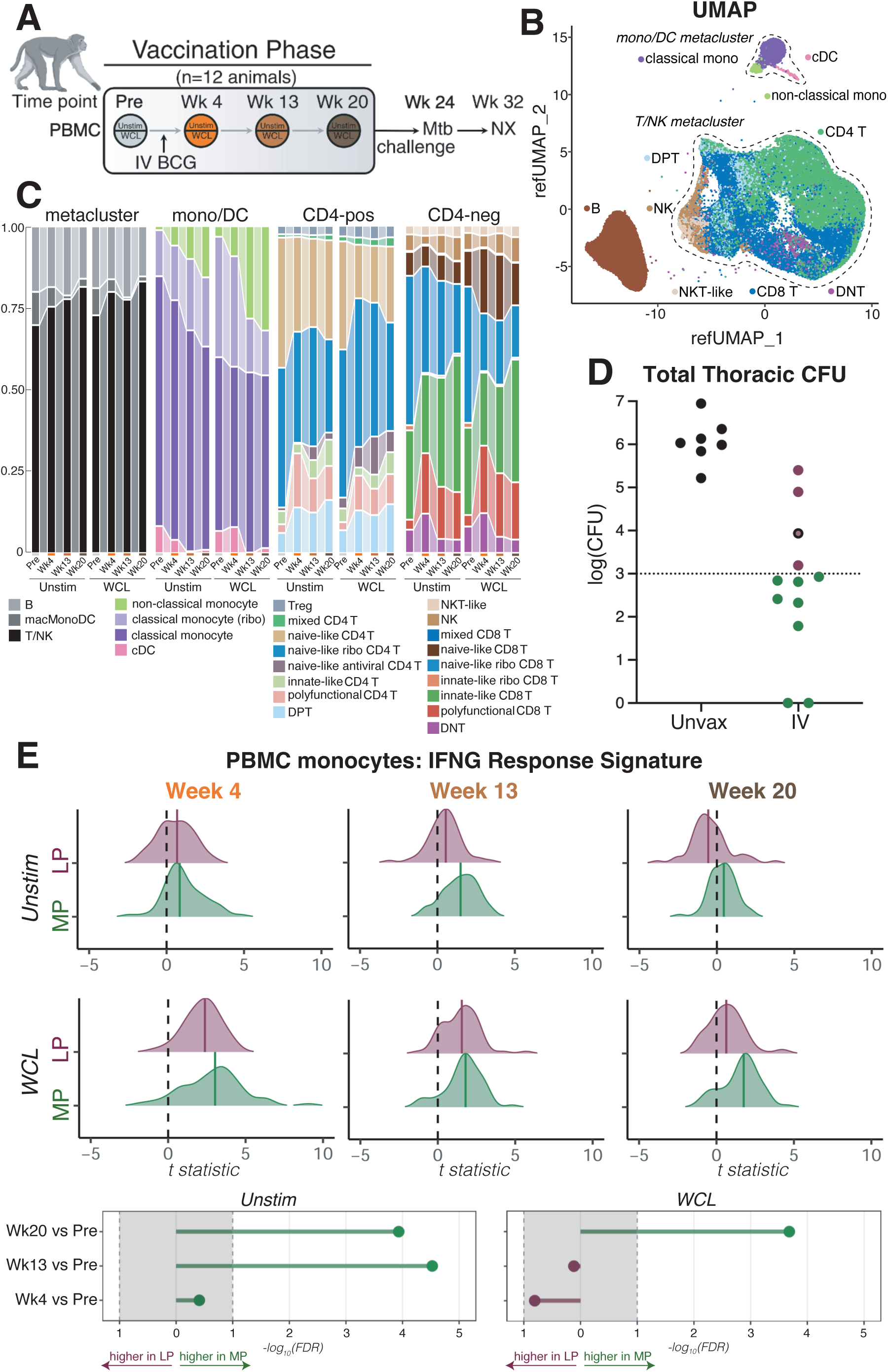
Independent cohort for correlating IFNG response signature with protection. **(A)** Experimental overview an independent cohort of macaques (n=12) vaccinated with IV BCG and challenged with Mtb Erdman 24 weeks later. Unstimulated (Unstim) and WCL-stimulated (WCL) PBMCs before (Pre) and after (Wk4, Wk13, Wk20) vaccination were collected and analyzed by CITE-Seq. **(B)** UMAP of PBMCs (*all cells*) from orthogonal dataset, reference-mapped to discovery vaccination phase PBMCs (Figure 1D). **(C)** Cell type composition (proportion of cell types) of PBMCs for *all cells* and for the mono/DC metacluster, CD4-positive T/NK metacluster, and CD4-negative T/NK metacluster; data are shown for each timepoint and stimulation condition (Unstim, WCL). **(D)** Total thoracic CFU (log_10_) measured 8 weeks after Mtb challenge in unvaccinated (unvax, n=7) and IV-BCG vaccinated (IV, n=12) macaques^13^. Dotted line represents categorical segregation of less protected (LP, >10^3^ CFU, purple) and more protected (MP, <10^3^ CFU, green) animals. Opaque symbol indicates IV BCG sample removed from downstream monocyte analysis due to cell thresholding. **(E)** Top, direction and strength of change in gene expression (*t* statistics) of the 61-gene IFNG response signature in pooled monocytes (classical and non-classical) from PBMC of less protected (LP) and more protected (MP) animals at Wk4, Wk13 and Wk20 (vs. Pre), with or without WCL stimulation. Solid lines are median *t* statistic; dotted line is zero enrichment. Bottom, statistical analysis (CAMERA competitive gene tests) of signature enrichment in monocytes from more protected (MP) vs. less protected (LP) at each timepoint (vs. Pre); dotted line represents significance threshold of FDR 0.1.

In the independent cohort of macaques, IV BCG conferred a 3.36-log reduction in mean CFU compared to unvaccinated controls (**Figure 7D**). Besides sterility, the definition of protection based on Mtb burden is arbitrary, however, the degree of control may be clinically meaningful. Prior to statistical analysis, we selected a cutoff of 10^3^ CFU to categorically partition IV BCG immunized macaques into “more protected” (<10^3^ CFU, n=8) and “less protected” (>10^3^ CFU, n=4) such that there would be adequate representation in each category. We then asked whether signatures defined in the discovery cohort associated with the degree of protection in the independent cohort or if they were general features of IV BCG immunization.

The IV BCG-specific double positive T (DPT) cell signature defined in PBMCs (Wk4 vs. Pre) of the discovery cohort (**Figure S4A**) was observed in DPT cells from all IV BCG immunized animals in the independent cohort, but with a larger effect size (*t* statistic) in more protected vs. less protected macaques (**Figure S15B**). Consistent with DPT cells responding to antigen, WCL-stimulated samples showed an increased DPT signature score compared to unstimulated samples. Furthermore, stimulation increased the difference in the median *t* statistic between more protected and less protected animals for DPT and polyfunctional CD4 T cells, indicating the genes in this signature may be associated with TB protection.

The 61-gene IFNG response signature observed in AMs and monocytes after vaccination in the primary dataset was evaluated in total monocytes from the orthogonal dataset, with or without WCL stimulation. In the absence of antigen stimulation, the signature was significantly enriched in monocytes in more protected vs. less protected animals at Wk13 and Wk20 (vs. Pre). This observation aligns with prior analysis suggesting a protracted activation state of BAL AMs after IV BCG (vs. naïve AMs), although IFNA response gene enrichment was greater than IFNG responses in the prior analysis^28^. Antigen stimulation positively shifted the 61-gene signature *t* statistics in all blood monocytes, but at Wk20 (vs. Pre) monocytes from more protected animals were significantly enriched compared to those from less protected macaques. These data substantiate the myeloid 61-gene IFNG response signature as a putative correlate of protection in blood.

## DISCUSSION

The ability of IV BCG immunization to prevent TB infection in macaques provides a pre-clinical model to assess immune correlates and mechanisms of protection in a naïve host that may inform TB vaccine design in humans. Here, we generated a longitudinal, multi-tissue rhesus macaque CITE-Seq atlas to compare the transcriptional profile of immune responses induced by IV and the benchmark ID BCG vaccine before and very early after TB challenge. The major findings include: 1) Definition of a 61-gene IFNG response signature in BAL AMs and blood monocytes with divergent expression between IV and ID BCG immunized macaques from Wk4 to Wk16 after vaccination; 2) Increased expression of the IFNG response signature and lower Mtb burden in lung from IV vs. ID immunized macaques at D11 post-challenge; 3) Association of the 61-gene IFNG response signature with protection in an orthogonal CITE-Seq dataset of blood monocytes from IV BCG-vaccinated, Mtb-challenged macaques; 4) Enhanced inferred communication between multiple *IFNG*-expressing T cell subsets and AMs in the airways of IV BCG-compared to ID BCG-vaccinated macaques; 5) Polyfunctional Th1/Th17-like DPT cells enriched for proliferative signatures were associated with enhanced protection following IV BCG in an independent blood analysis.

T cells are required for protection against primary TB and TB reinfection, and for IV BCG-elicited protection^14,26,27,46^. Prior studies showed that higher frequencies of Th1/Th17-cytokine producing CD4 T cells and NK cells in the airway (BAL) correlated with protection after IV BCG^12^; however, there were no clear blood correlates of protection defined by a multi-dimensional cellular and humoral immune analysis^12^. Here, by extending the analyses using CITE-Seq, we show that while most T cells in PBMC are naïve- or innate-like, relative proportions of polyfunctional CD4 and CD8 T cells significantly increased following IV compared to ID BCG vaccination. Further, we observed an expanded population of CD4+CD8ɑ+ DPT cells in blood in IV BCG that was enriched (compared to ID BCG DPT cells) for Th1/Th17 modules previously described in BAL T cells after BCG vaccination^11,28^. DPT cells from IV BCG vaccinated macaques displayed the highest level of DE among all cell types tested at Wk4 and Wk16 (**Figure 1L**) with genes enriched (ORA and GSEA) for pathways of activation and proliferation. Since DPT cells express both CD4 and CD8ɑ, loss of this population might have contributed to significantly reduced protection following depletion of CD4 or CD8ɑ-expressing cells after IV BCG^14^. In cynomolgus macaques, scRNA analysis of lung tissue after primary infection showed that a cluster of *CD4+, CD8A/B+* Th1/Th17-like cytotoxic T cells were associated with lower-burden granulomas^24^. Lastly, in humans, the percentage of DPT cells in leukocytes was reduced in active TB patients and inversely correlated with TB disease severity^47^.

The CITE-Seq analysis in this study also identified key transcriptional differences in myeloid cells between IV and ID BCG macaques that evolved over time after vaccination and were recapitulated shortly after TB challenge. In humans, myeloid reprogramming after ID BCG results in increased cytokine responses to heterologous innate stimuli and enhanced protection against viral infection, a process termed trained immunity^48–50^. In mice, systemic (IV) but not peripheral (subcutaneous) BCG administration protects against homologous and heterologous challenges, indicating that the IV route is a more effective inducer of enhanced myeloid function in this model^18–20^. In macaques, enhanced innate cytokine production and histone modifications consistent with innate training were reported in PBMCs two weeks after IV BCG vaccination^51^. However, prior reports show the durability of trained immunity after vaccination may be limited^50,52^. The data presented here show IV BCG elicits sustained IFNG response signaling in blood monocytes and BAL macrophages compared to ID BCG before and after TB challenge. These data extend our prior analyses that reported a late IFNA/IFNG response signature in BAL AMs after IV BCG but did not assess blood or an association with protection^28^. We hypothesize that increased and sustained IFNG responsiveness in AMs may have largely mediated the reduction in lung Mtb CFU observed here 11 days after challenge in IV BCG macaques as AMs are the initial cells in BAL to encounter Mtb^22,23^. This is further supported by post-challenge barcode analysis in macaques from prior studies demonstrating IV BCG restricts the number of Mtb bacilli that establish infection even after T cell depletion^14^, whereas ID BCG fails to do so. These data are consistent with a recent report in a mouse strain with impaired T cell trafficking showing that early TB control after BCG immunization was mediated by an innate response, independent of T cell memory responses^53^.

ORA and GSEA of BAL AMs indicated significant IFNG *and* IFNA response signaling (IV vs. ID) between Wk4 and Wk16 after BCG. Similarly, IFNA responsiveness in monocytes/AMs from the lung after primary MTb challenge^25,27^ and after MtbΔSigH or IV BCG immunization^10,28^ has been reported based on enrichment of downstream gene sets as done here and sender-agnostic cell-cell communication analyses, rather than by detection of the IFNA ligand itself. Indeed, *IFNA* transcript detection is limited due to sparse RNA read recovery and mapping ambiguity of the highly homologous macaque *IFNA* locus in short-read scRNA-seq used here, which in the future may be overcome by targeted quantification using long-read sequencing. Further, IFNA and IFNG response gene sets (pathways) share substantial transcriptional overlap. Here, by partitioning the Hallmark IFNA response signatures into IFNG-shared and IFNA unique genes, our analyses suggest that some IFNA response enrichments after BCG immunization may instead reflect a broader IFN-responsive program associated with strong IFNG activity. We therefore focused subsequent analyses on the more robust IFNG response program to define a discrete IFNG response gene module linked to protection. This approach is supported by the established role of IFNG in trained innate immunity and T cell mediated protection against TB, as well as by concordant protein-level IFNG production by antigen-stimulated PBMCs and BAL after IV BCG in our study.

Multiple mechanisms could account for sustained IFN responses in myeloid cells following IV BCG compared to ID BCG. Systemic delivery and a higher dose of IV BCG compared to ID BCG leads to greater persistence of antigen in multiple tissues, providing a durable stimulus for myeloid and T cells^11^. Additionally, systemic IV delivery of BCG has been shown to facilitate “central” trained immunity via hematopoietic stem cell (HSC) reprogramming in mice^17^. This could be partially due to PRR-mediated innate cytokine signals in the blood or bone marrow early post-vaccination. Indeed, a previous study using bulk RNA-seq to profile a dose-ranging cohort of IV BCG-vaccinated macaques showed that an early (D2) antiviral, Type I IFN response signature from whole blood was predictive of peak T cell responses in BAL after vaccination and TB outcome following a 12-week challenge phase^29^. It remains unclear which cells drive this Type I IFN response transcriptional signature as the CITE-Seq dataset here did not profile macaques immediately post-vaccination. In addition to central trained immunity, “peripheral” training of mature myeloid cells can occur locally in tissues through IFNG production by CD4 T cells, providing heterologous protection against respiratory viruses in the lungs of mice after IV BCG^18–20^. Here, CellChat analysis inferred that multiple IFNG-producing T cell subsets, including polyfunctional T cells (CD4, CD8, DPT, anti-viral CD8) and MAIT-like T cells were communicating with AMs in the BAL after IV BCG. In contrast, IFNG signaling to AMs was limited to polyfunctional CD4 T cells after ID BCG. Thus, these data confirm and extend previous findings that IV BCG elicits a higher magnitude of antigen-specific T cell responses compared to ID BCG by revealing greater breadth of IFNG-producing T cell subsets in the BAL following IV vaccination in the lung. Such diversity in CD4 and CD8 T cell-derived IFNG provides compensatory mechanisms to sustain IFNG responsiveness in AMs and may explain why CD4- or CD8-depleted, IV BCG-vaccinated macaques retained enhanced ability to restrict early Mtb establishment after infection^14^.

An ongoing objective for TB vaccine development is to define predictive blood signatures of protection. Studies in humans have primarily defined correlates of risk for developing TB disease^54^. Our previous multimodal analysis of dose-escalated IV BCG-vaccinated macaques failed to identify predictive blood correlates of protection, though this work primarily utilized flow cytometry and systems serology data^12,15^. The more comprehensive and unbiased single cell CITE-Seq approach presented here harnessed strong vaccine-induced gene expression changes in BAL and a matched PBMC dataset to define a myeloid IFNG response signature as a candidate blood correlate of protection. The 61-gene IFNG response signature was defined in BAL AMs by comparing differential gene expression in paired longitudinal samples collected before and after ID and IV BCG vaccination and was recapitulated in matched monocyte samples from PBMCs. The signature was enriched in AMs early post-challenge in IV BCG-immunized animals with reduced Mtb burdens compared to ID BCG and unvaccinated macaques, indicating the signature reflects protective immunity in the airway and lung. Although the 61-gene signature was defined by contrasting IV vs. ID samples, it was also enriched in blood monocytes from more protected vs. less protected macaques, all of which were immunized with IV BCG (independent cohort). Hence, this IFNG response signature is the first candidate of a cell-type specific, BCG route- and dose-independent blood correlate of protection. It will be important to determine in future pre-clinical and human vaccine studies whether this signature applies to other live attenuated vaccines or alternative subunit approaches. However, this work suggests that studies defining vaccine correlates should evaluate both peak and memory timepoints given the IFNG response signature genes here were upregulated from Wk4 to Wk16 post-vaccination. Based on its predictive role for vaccine elicited protection and its potential role for early control of TB in the lung, the IFNG response signature could be useful in evaluating and prioritizing pre-clinical vaccine candidates.

The 61-gene myeloid signature includes genes involved in antigen processing and presentation, T cell co-stimulation and adhesion, T cell recruitment, and immunoregulation that collectively may contribute to improved control of Mtb by AMs upon infection. Some genes within the signature such as CXCL9/10/11 were identified in IFN-responsive macrophages in the BAL after IV BCG and from low-burden granulomas after primary infection^24,28^. However, in other contexts of macaque and human TB progression, expression of CXCL9/10/11 and IFN signaling in myeloid cells appeared driven by antigen burden^25,27,55^. Similarly, genes involved in inflammation and immunoregulation such as IDO1 and CD274 which were upregulated after IV BCG may be interpreted differently with respect to protection depending on the presence of ongoing infection^25,27,55^. Overall, these data highlight differences in protective vaccine-mediated immune responses versus less protective primary immune responses despite some pathways being enriched in both contexts.

The work presented here has several limitations. IV BCG induces epigenetic changes in bone marrow HSCs and mature myeloid cells in mice, which is thought to be a primary mechanism of trained immunity^17,50,51^. Epigenetic analysis in blood and bone marrow was beyond the scope of this study and will be performed in future studies. Additionally, we did not perform *in vitro* growth inhibition assays using myeloid cells from IV vs. ID immune macaques, although the early reduction in Mtb burden in the lung and lymph nodes after IV BCG vaccination provides evidence for enhanced function. The conclusions on T cell-AM communication are inferred from transcriptional ligand-receptor expression as we did not directly measure T cell-AM crosstalk. However, IFNG protein in response to antigen stimulation was higher in blood and BAL after IV compared to ID BCG by both flow cytometry and Luminex assays. Still, it is possible that the IFNG-responsive pathways are enriched through other cellular interactions, such as IFNA, which is not easily quantified transcriptionally in macaques. Finally, while this study analyzed the vaccine responses from 9 animals before and after IV or ID BCG, the post-challenge cohort was limited to 3 animals per group. Future work will focus on understanding mechanisms of protection using larger cohorts.

This study provides a framework for understanding high-level vaccine-induced protection against TB, including prevention of infection as well as disease. We propose that integrated innate and adaptive immunity in the lung is critical for early and sustained vaccine-mediated protection against TB. IV BCG elicits a high magnitude of CD4 and other T cell types to produce IFNG that induces and sustains IFNG response pathways in myeloid cells in blood and at the site of infection. This study provides hypothesis-generating data that early control of TB in macaques after IV BCG may be achieved by activated AMs with CD4 T and CD8ɑ-expressing cells required to sustain protection in the lung^56^. This model provides a roadmap for improving ongoing TB vaccine efforts.

## ACKNOWLEDGEMENTS

This research was supported by the Intramural Research Program of the National Institutes of Health (NIH). The contributions of the NIH author(s) are considered Works of the United States Government. The findings and conclusions presented in this paper are those of the author(s) and do not necessarily reflect the views of the NIH or the U.S. Department of Health and Human Services. JDD is currently supported by NIH Medical Scientist Training Program Training Grant T32GM136651. Data in Figure S11A was in part supported by the National Institutes of Health [IMPAc-TB (HI-IMPACT)] 75N93019C00071. The following reagent was obtained through BEI Resources, NIAID, NIH: *Mycobacterium tuberculosis*, Strain H37Rv, Whole Cell Lysate, NR-14822. We thank members of the VRC Flow Cytometry Core for BSL2 cell sorting and support; VRC Translational Research Program for animal program assistance; TB Imaging Program and Comparative Medicine Branch of NIAID for animal study support. This work utilized the computational resources of the NIH HPC Biowulf (https://hpc.nih.gov) and Skyline (https://skyline.niaid.nih.gov/) clusters. Illustrations in Figures 1A and 4A were created with Biorender.com.

## AUTHOR CONTRIBUTIONS

Conceptualization: RAS, PAD, JDD

Investigation: JDD, MAR, ANB, CCL, FL, ARH, LEV, AMW, MC, JL, MSM, MCC, FH, TF, JLG

Formal analysis: PLM, JDD, MAR, PAD

Software: PLM, MAR, JDD

Writing-original draft: JDD, PLM, MAR, PAD, RAS

Writing-review and editing: JDD, PLM, MAR, PAD, RAS, SMF, CS Supervision: LEV, ID, CS, JAF, SMF, DCD, MR, PAD, RAS

## METHODS

### Animals and handling

3-5 year old Indian-origin rhesus macaques (n=18) were used in the “signature discovery cohort” (**Table S1**); all were male based on animal availability. Study design and procedures were approved by the NIH NIAID and VRC Animal Care and Use Committees (ACUC). Macaques were housed and cared for in accordance with local, state, federal, and institute policies in facilities accredited by the American Association for Accreditation of Laboratory Animal Care (AAALAC), under standards established in the Animal Welfare Act and the Guide for the Care and Use of Laboratory Animals. Macaques were provided intellectual and physical enrichment and monitored for physical health, food consumption, body weight, temperature, complete blood counts, and serum chemistries. Methodology related to macaques (n=12) from which cryopreserved samples were analyzed in the “independent cohort” (**Table S1**) was reported previously^14^.

### BCG Vaccination

Macaques were vaccinated with BCG Danish 1331 that had been expanded and cryopreserved. IV BCG (2.8–8.1 x 10^7^ CFU) or ID BCG (3.9–6.2 x 10^5^ CFU) was administered as described previously^11^. BCG dose for each animal is in **Table S1**.

### Tissue Processing

Peripheral blood and BAL fluid collected before and after (Wk4, Wk16) vaccination was processed as previously described^11^. Peripheral blood, BAL fluid, and lung tissue were collected 11 days post-challenge. Infected lung tissue was manually minced and enzymatically digested in RPMI containing Liberase TL (100 ug/mL), hyaluronidase (240 ug/mL), and DNAse (50 U/mL) from Sigma for 40min in a shaking incubator (150 rpm, 37°C) and then FBS was added to a 10% final concentration. Lung tissue was passed through GentleMacs 70Um filters (Miltenyi), and an aliquot was removed for CFU plating. Cells were centrifuged (10min, 1500 rpm), resuspended in 10mL ACK lysis buffer (4min), washed in PBS twice, and resuspended in 10mL RPMI containing 10% FBS. All samples were cryopreserved in FBS/10% DMSO in a -80°C freezer.

### Flow Cytometry

Cryopreserved peripheral blood and BAL samples were thawed, rested, stained, and analyzed in batches as described previously^11^. Antibodies used are listed in **Table S4**. Samples were analyzed on modified BD FACSymphony machines and analyzed using FlowJo (v.10.10.0).

### Mtb challenge and burdens

Prior to challenge, macaques were transferred to a BSL-3 animal facility and housed individually in biocontainment cages approved for the containment of *Mtb*. Macaques were challenged 5–6 months after BCG vaccination with 16– 37 CFU of Mtb Erdman mCherry in a 2 ml volume in the right lower lung lobe using a bronchoscope (**Table S1**). Macaques were humanely euthanized 11 days post challenge. Tissues plated for Mtb burden quantification included: lung lobes (right upper, right lower, left lower), spleen, liver, blood, BAL and extra-pulmonary lymph nodes (axillary, inguinal), pulmonary lymph nodes (hilar, carinal). A portion of each homogenized tissue was plated on 7H11 agar (Difco) and incubated at 37° C for three weeks. Only lung tissue and lung lymph nodes had detectable Mtb. CFUs were back-calculated based on the total amount (grams) of tissue processed.

### Mtb Barcode Analysis

Mtb genomic DNA was purified from CFUs of infected tissues as described^57^. Genomic DNA was subjected to transposon-mediated fragmentation (Nextera XT, Illumina) and paired end sequencing (150 bp) at SeqCenter (Pittsburgh, PA) and BioMicroCenter (MIT). Fastq reads were passed to a custom script to quantitate unique barcodes associated with the Mtb library as described^58^. Briefly, all singleton barcodes were removed as sequencing artifacts and the remainder were quantified as true barcodes in each tissue. Unique barcodes summed across all tissues per animal reflects the number of Mtb bacilli that established infection. Detailed descriptions for whole genome sequencing based barcode detection scripts are at: https://github.com/Fortune-Lab/Mtb-WGS-barcoding

### CITE-Seq Library Generation

Cryopreserved vaccination phase PBMC and BAL cells were thawed and rested in R10 (RPMI 1640 + 10% heat-inactivated FBS) for 8 hours at 37℃. Cells were then plated in 96-well plates at 2×10^6^ cells/well and stimulated with 20 ug/mL Mtb H37Rv whole-cell lysate (WCL, BEI Resources) for 8 hours at 37℃. Following stimulation, supernatant was collected for Luminex analysis (NHP XL Cytokine Luminex Performance Premixed Kit, Bio-Techne, PN: FCSTM21-24, per manufacturer’s instructions). Samples were resuspended in PBS + benzonase nuclease (Sigma, 50 U/mL) and replated at 5×10^5^ (BAL) or 1×10^6^ (PBMC) cells/well. Cells were washed in PBS, then incubated with human FcR block for 10 minutes. Cells were washed with PBS/ 10% BSA (FACs buffer), then incubated with hashtag oligonucleotides (Biolegend) for 20 minutes. Samples were washed with FACs buffer and pooled to minimize batch effects and improve throughput. Pooled samples were stained for 20 minutes with a cocktail of human CITE-Seq antibodies (Biolegend, **Table S4**) plus anti-CD45 FITC optimized for rhesus macaques. The samples were washed, resuspended in sort buffer (PBS/2% FBS), and filtered through 35 μm filters. SYTOX Red Live-Dead stain (Thermo) was added just prior to sorting (1:1000) to label dead and dying cells. Samples were sorted for live CD45+ cells using a FACSAria III cell sorter (BD Biosciences). Sorted cells were loaded at a concentration estimated to produce under a 20% doublet rate into a 10x Genomics Chromium X. The 10x Genomics 5’ V2 library preparation kit and protocol were used to generate RNA and protein barcode libraries. Quantification of initial cDNA amplification as well as final library concentrations were performed using the Agilent Bioanalyzer High Sensitivity DNA assay. Gene expression and CITE-Seq antibody derived tag cDNA libraries were sequenced with either S4 flow cells on an Illumina NovaSeq 6000 with S4 flow cells or on an Illumina NovaSeq X with 10B flow cells.

Challenge phase PBMC, BAL, and lung cells were processed similarly to pre-challenge samples except infected samples were not stimulated with Mtb WCL prior to staining and sorting. Infected samples were processed under BSL3 conditions with 10x Chromium Controller and sorted for live CD45+ cells using a FACSAria III cell sorter (BD Biosciences) at BSL-3.

### Single-cell CITE-seq data processing and integration

#### CITE-seq library multiplexing

Samples were multiplexed using unique barcodes via hashtag oligonucleotides (HTO) attached to antibody-derived tags (ADT) to enable sample multiplexing (n=2-6 samples per GEM well).

#### Cell counts and normalization

Cellranger count (v.7.2, 10x Genomics) was run to generate gene and protein expression counts from raw fastq files, aligning to the rhesus macaque genome (Mmul_10, protein-coding, Ensembl release 101, GCF_003339765.1). For the vaccine phase PBMC and BAL analyses, 518,718 droplets were collected from 21 subjects across 3 timepoints (Pre, Wk4, Wk16) and three conditions (ID BCG, IV BCG, unvaccinated) and 4 batches. Within each GEM well, gene expression was normalized by removing ambient RNA (filtered expression matrix) using SoupX v1.6.2^32^ and the autoEstCont and adjustCounts function, and normalized and denoised protein expression (ADT raw expression matrix) using dsb^33^ and the DSBNormalizeProtein function, with isotype controls (mIgG1 K, mIgG2b K, mIgG2a K, Armenian Hamster IgG).

#### Demultiplexing

Samples were demultiplexed by first applying Seurat::FindNeighbors and Seurat::FindClusters (k.param=20) to the hashtag ADT data, manually assigning clusters of cells to animal IDs, and then removing unclassified singlets and heterotypic doublets (Seurat v5.0.2)^59^

#### Quality control

After demultiplexing and doublet removal, 423,759 singlets were identified. To remove low-quality cells, QC was performed, retaining singlets based on # of features ( 250 < n < 5,000) and mitochondrial genes composition (< 20%, gene list: *ND2, ND3, ND4, ND4L, ND5, COX1, COX2, COX3, ATP6, ATP8,* and *CYTB*). This filtering retained 419,395 high-quality singlets (n=198,453 PBMC; n=220,932 BAL) in the vaccine phase and 88,109 high-quality singlets (n=26,576 BAL, n=39,657 LUNG) in the challenge phase.

#### Seurat and Harmony clustering workflow

Singlets were sequentially clustered using Seurat. In each case, a standard Seurat v5 method was used to normalize RNA counts and UMIs, converting counts to natural log units with a scale factor of 10,000 with NormalizeData, z-scoring the RNA count data using ScaleData, identifying the top 2,000 highly variable features using FindVariableFeatures (method=“vst”), and applying principal components analysis using RunPCA. Prior to cell clustering, the effect of separate genome sequencing runs (tenXrun) was removed using RunHarmony from the harmony package (v1.2.0)^60^, a shared nearest-neighbor graph was constructed using FindNeighbors (k.param=20, dims=1:30), clusters of related cells were identified using FindClusters (at varying resolutions; Louvain method^61^, and a reduced dimension map of embedding was generated using the Uniform Manifold Approximation and Projection (UMAP) approach implemented in RunUMAP (dims=1:30).

### Clustering and Cell Type Assignment

*Vaccine Phase.* For the PBMC analysis, an initial Seurat clustering analysis was performed to identify 30 unsupervised clusters, removing 3 clusters that were low-quality, resulting in n=192,834 cells. For the BAL analysis, an initial Seurat clustering analysis was performed to identify 26 unsupervised clusters, removing 2 clusters that were low-quality, resulting in n=216,446 cells. After removal of low-quality clusters, clustree (v. 0.5.1)^62^ was used for visualization, and gene expression markers and ADT markers were used to optimize clustering resolution and determine cell type identity, identifying n=18 PBMC and n=20 BAL clusters which were then manually pooled into “metaclusters” based on coarse cell type lineages (T/NK, mac/mono/DC, B cell, or basphil). For the BAL, the smallest cluster (#19, too few cells for downstream analysis) was removed, and cells in a mixed T cell / macrophage cluster were reassigned to their respective metaclusters based on UMAP dimension 1 coordinates.

Within the macMonoDC clusters (cell counts PBMC n=5,876, BAL n=68,321), after re-clustering, low-quality and low-frequency clusters (<100 total cells) were removed, resulting in n=5,719 and n=62,909 cells in PBMC and BAL respectively. Unvaccinated samples contained low or highly variable cell counts and were removed for downstream gene expression and gene set enrichment analysis. The resulting cell clusters were annotated as macMonoDC “fine” cell types. For the BAL, 4,684 CD2-high cells that were classified as alveolar macrophages were identified and removed, and 40,976 CD2-negative alveolar macrophages were retained for downstream analysis.

Within the T/NK clusters (PBMC n=157,268, BAL n=140,266), linear gating was performed based on CD4 and CD8 protein (ADT) expression to generate CD4+ and CD4− T/NK metaclusters. Within each of these metaclusters, cells were re-clustered and low-quality and low-frequency clusters were removed resulting in a final CD4+ (PBMC n=86,547; BAL n=38,678) and CD4− (PBMC n=70,696; BAL n=101,588) set of high QC cells. These cell clusters were also annotated as T/NK “fine” cell types, and joined with the macMonoDC annotations, the B cell annotations (B_1, B_2, B_3 for PBMC [n=5,876], and a single B cell cluster for BAL [n=4,739]), and basophils (n=1,702) which were present in BAL only to an “all cells” object. This resulted in 186,952 (PBMC) and 201,609 (BAL) high-quality cells, across a total of n=53 fine clusters, which were pooled into n=25 intermediate clusters that were used in downstream cell type frequency and gene expression analysis (DEGs, GSEA, etc.) and further pooled into n=10 broad clusters (Fig 1D, 1E). Cluster-specific gene expression markers were identified at each resolution through differential expression analysis (wilcox.text, presto v1.0.0)^60^. The DE markers as well as canonical markers from the literature were used to assign cell type identities and states to each of the clusters.

#### Challenge Phase

The same clustering and cell type assignment process was applied separately to challenge phase BAL and lung data, resulting in 25,710 high-quality BAL cells and 39,023 lung cells.

### Calculation of pseudobulk sums

To generate sample-level RNA counts (“pseudobulk”), cells were assigned to sample-specific groupings defined by the relevant combination of subject ID, timepoint (vaccine phase), sequencing run, and experimental batch, applicable to each phase of the study. An indicator matrix was constructed using the Matrix::sparse.model.matrix function from the Matrix package (v1.6-5), https://dx.doi.org/10.32614/CRAN.package.Matrix, formula = ∼ 0 + sample). The gene-by-cell RNA count matrix was multiplied by this indicator matrix to obtain a pseudobulk count matrix, summing raw RNA counts across all cells belonging to each sample.

### Multidimensional scaling of pseudobulk samples across cells

To examine the global patterns of gene expression across pseudobulk samples, filtered and normalized pseudobulk counts were first transformed to log-expression values using voom (limma). For visualization, nuisance effects were removed from the voom expression matrix using removeBatchEffect (limma), preserving the biological group effects of interest. For the vaccine phase, effects of experimental block, age, and weight were removed; for challenge phase data, effects of 10x sequencing run were removed. Pairwise Euclidean distances between samples were calculated from the corrected expression matrix, and classical multidimensional scaling (MDS, principal coordinates analysis) was performed using stats::cmdscale^63^. Classical MDS on Euclidean distance matrices yields the same sample configuration as PCA.

### Cell type composition

For each tissue compartment and cell type resolution, cell type frequencies were calculated separately for each biological sample. Populations included broad metaclusters (e.g., TNK, macMonoDC, B, basophil), TNK subpopulations (e.g., DPT, DNT, pooled CD4 T, pooled CD8 T, MAIT-like), and intermediate cell-type clusters. Differences in cell type composition were tested using generalized linear models (quasibinomial, allowing for overdispersion) applied to the observed cell counts (cbind(n, total -n)). Comparisons between vaccine groups included vax_group as the predictor. Longitudinal comparisons within vaccine groups, across timepoints, additionally included subject_ID as a fixed effect. FDR adjustment (BH, Benjamini & Hochberg)^64^ was applied across all cell type × contrast tests within each tissue/resolution analysis, and statistical significance was defined as *p.adj* < 0.05.

### Modeling gene expression across tissues and timepoints

To model the effect of vaccination route and time point on gene expression levels, for each cell type [intermediate cluster resolution] the following linear models using limma and voom^37^ were fit to model the mean-variance relationship of log counts and obtain precision weights for linear modeling. For the vaccine phase, voomLmFit (edgeR v4.6.3)^65^ was used with sample quality weights and and duplicateCorrelation (limma package) applied to subject ID, to account for repeated measures of the same animals across the time course. For the challenge phase, voomWithQualityWeights (limma v3.64.3)^66^) [challenge phase] was used to account for heterogeneity in quality across sample and observation level counts. We restricted differential gene expression analysis to cell types where we had at least 30 cells per vaccine group and timepoint required for the contrast [vaccine phase] or 10 cells per individual sample [challenge phase]. For each celltype, we restricted the number of genes tested using filterByExpr() from the edgeR package, with the following standard parameters (min.prop = 0.7, min.count = 10, min.total.count = 15). Animal age and weight values were z-scored across individuals using the scale() function in R (R base, v. 4.5.1), separately for the vaccine phase and challenge phase.

For the vaccine phase data, we first modeled the pseudobulk counts by sample, accounting for fixed effects of batch (experimental block and sample run), age, and weight. For the challenge phase data, we modeled the pseudobulk counts by sample, accounting for fixed effects of sample run (tenXrun) and age.

Vaccine phase: E ∼ 0 + timepoint_vax_group + age + weight + block

Challenge phase: E ∼ 0 + vax_group + age + tenXrun

where block is the combined factor of tenXrun_batch.

For the vaccine phase analysis, up to nine prespecified contrasts were evaluated for each cell type, using contrasts.fit^67^ conditional on sufficient cell representation:

1. Week 4 versus baseline within ID
2. Week 16 versus baseline within ID
3. Week 16 versus Week 4 within ID
4. Week 4 versus baseline within IV
5. Week 16 versus baseline within IV
6. Week16 versus Week 4 within IV
7. IV vs ID difference in baseline to Week 4 change
8. IV vs ID difference in baseline to Week 16 change
9. IV vs ID difference in Wk4 to Week 16 change

Moderated t-statistics were obtained using empirical Bayes (limma:: eBayes), and for each contrast, gene-level P-values were adjusted separately using the BH procedure. Genes with FDR < 0.10 were considered differentially expressed.

### Cross-species ortholog identification

To enable human gene set-based functional enrichment analyses of the macaque gene expression results, a macaque-to-human ortholog mapping resource was first generated for all profiled macaque gene features (n=21,369), including features represented by Ensembl gene identifiers. Ortholog mappings were obtained using orthogene convert_orthologs() from the orthogene package (v1.14.0)^68^ and gorth() from the gprofiler2 package (0.2.3)^69^. Mapping was performed in both macaque-to-human and human-to-macaque directions, and unique mapping pairs were combined to characterize one-to-one and multi-mapping orthology relationships. Reference annotations were obtained from Ensembl release 113; rhesus macaque assembly Mmul_10; and human assembly GRCh38.p14. Macaque gene identifiers were classified as having no known human ortholog (n=4,300), mapping one-to-one (n=16,217), one-to-many (n=155), many-to-one (n=452) or many-to-many (n=245).

### Gene set overrepresentation analysis (ORA)

We constructed 2×2 contingency tables based on vaccine phase DEG lists and HALLMARK PATHWAYS (n=50) (msigdbr v7.5.1, https://cran.r-project.org/web/packages/msigdbr/index.html). We use the fisher.test() function from the stats package (v.4.3.2, https://www.R-project.org/; options: alternative = “greater”), to assess the independence of a feature in a specific celltype/timepoint DEG set relative to the universe of genes tested for DE in the corresponding tissue (n=12,325 tested in PBMC or n=14,328 tested in BAL), where genes were considered DE at BH-adjusted p<0.1 and |logFC| > 0.25.

Fisher’s exact-test P-values were calculated from the unmodified contingency tables. Separately, we used a Haldane-Anscombe correction^70,71^ to obtain stabilized log odds ratio estimates and approximate confidence intervals.

logOR = log(((a+0.5)*(d+0.5))/((b+0.5)*(c+0.5)))

se = sqrt(1/(a+0.5) + 1/(b+0.5) + 1/(c+0.5) + 1/(d+0.5))

Fisher test *P*-values across Hallmark pathways were adjusted using the BH procedure within each ORA.

### Gene set enrichment analysis (GSEA)

To identify relevant functional pathways associated with vaccination route and timepoint, for each cell type and DE contrast, we ranked genes using a signed significance statistic, -log10(*P*) × sign(logFC). Gene set enrichment analysis was then performed using fgseaMultilevel from the fgsea package (v1.28.0)^72^ against the macaque-converted Hallmark gene sets.

### Cell-cell interaction analysis

We used CellChat (v2.1.2)^73^ to infer ligand-receptor interactions across cell types (intermediate resolution) by integrating our single-cell gene expression data with the human database, CellChatDB v1, which contains 291 annotated pathways; after restricting to protein-mediated interactions, 253 pathways were retained for analysis. We restricted our analysis to cell types (n=17) with 1,000 cells or more, excluding pDC, Treg, basophils and DNT because of low representation. We first randomly downsampled the BAL to 19,000 cells per timepoint-by-vax combination. We annotated overexpressed signaling genes and ligand-receptor interactions in each cell type using identifyOverExpressedGenes and identifyOverExpressedInteractions, respectively.

We then computed the communication strength using computeCommunProb with and used the “triMean” method to calculate the average gene expression of ligands and receptors for each cell group. Communication probabilities were inferred using computeCommunProbPathway(type=“triMean”, population.size=FALSE), and generated an aggregated network, counting the number of links, or summation of communication probabilities, using aggregateNet. After running the individual analyses for each timepoint_vax group, we performed a pairwise comparison of cellChat across the following timepoints and conditions:

1. Wk4 vs. Pre within ID
2. Wk16 vs. Pre within ID
3. Wk16 vs. Wk4 within ID
4. Wk4 vs. Pre within IV
5. Wk16 vs. Pre within IV
6. Wk16 vs. Wk4 within IV
7. IV vs ID at Pre
8. IV vs ID at Wk4
9. IV vs ID at Wk16

For select cell-cell communication analyses and visualizations, we then focused on just the interactions between alveolar macrophages and T cell subsets: polyfunctional CD4 T, polyfunctional CD8 T, polyfunctional antiviral CD8 T, DPT, and MAIT-like.

We used compareInteractions to compare the different number of inferred interactions between each pairwise set of data. We used rankNet to identify differences in pathway-level information flow across conditions using CellChat’s paired Wilcoxon test (*P* < 0.05). We then performed differential expression analysis for each pairwise set of conditions, within each cell group, for signaling-associated genes within each cell group using identifyOverExpressedGenes (fraction of cells expressing gene in cluster > 0.1; logFC > 0.05, p < 0.05). Differentially expressed signaling genes were then mapped to ligand-receptor interactions using netMappingDEG.

### Definition and testing of single-cell signature scores

We defined a 61-gene divergent IFNG response signature as the intersection of the leading edge genes (LEGs) from the Hallmark INTERFERON_GAMMA_RESPONSE GSEA for the Wk16 vs. Wk4 contrast within the IV and ID vaccine groups in vaccine phase BAL alveolar macrophages. The shared LEGs reflected opposing longitudinal responses, with higher expression at Wk16 vs Wk4 in the IV group, and lower expression at Wk16 vs Wk4 in the ID group. We visualized the distributions of limma/voom *t* statistics for these genes using ridgeline plots to qualitatively compare the direction and strength of differential expression of the signature genes across vaccine phase BAL alveolar macrophages and blood classical monocytes, as well as in the challenge phase BAL and lung alveolar macrophages.

For the vaccine phase, to quantitatively compare the signature scores on a single cell level across vaccine groups at each timepoint, we generated single-cell module scores and used lme4 (v1.1-36)^74^ and emmeans (v1.10.1, https://CRAN.R-project.org/package=emmeans) for testing and visualization. Module scores were calculated separately in vaccine phase BAL alveolar macrophages and blood classical monocytes using AddModuleScore_UCell from the UCell package (v2.12.0)^75^.

To adjust the module scores, and remove nuisance effects, we first fit a linear mixed model with covariates time point and vaccine group, and nuisance variables (age, weight, and block, which is batch_tenXrun), as well as a random effect for subject ID and residual error:

UCell_score ∼ 0 + group + block + age + weight + (1 | subject)

Where group denotes the six timepoint-by-vaccine combinations, and block = batch ×10x_run.

We then calculated adjusted module scores for each cell are given by:

adjusted_Score = observed_UCell_score − block effect − age effect − weight effect − subject random intercept, while retaining the estimated timepoint-by-vaccine group effect.

Group estimated marginal means and prespecified contrasts were obtained from the fitted mixed effects model using emmeans (model: ∼ group; lmer.df = “satterthwaite”). The adjusted single-cell scores were used for visualization and statistical inference was based on the fixed effects model (within contrast groups, FDR adjust=“BH”).

Also, a 100-gene DPT signature was defined from vaccine phase blood DPT cells using the Wk4 IV-versus-Pre contrast. IV-specific upregulated genes were ranked by BH-adjusted P-value, with log2FC used to break ties, and the top 100 genes were retained. Candidate genes were required to be upregulated in IV animals (log_2FC>0.25, BH-adjusted P<0.10) and were excluded if they also positively associated with the corresponding ID contrast at a relaxed threshold (BH-adjusted P<0.40, log_2FC>0.25).

To test predefined gene signatures while accounting for inter-gene correlation, competitive gene-set testing was performed using CAMERA/CAMERA-PR from the limma package^38^. The divergent IFNG response signature was tested across relevant vaccine and challenge phase differential-expression contrasts and the DPT signature was tested across vaccine phase contrasts. CAMERA was used for vaccine phase analyses and CAMERA-PR for challenge phase analyses, using moderated t-statistics from the corresponding DE contrasts as gene-level statistics (inter-gene correlation set to 0.01).

#### AI-assisted code development

ChatGPT (GPT-5.6 Sol, OpenAI) and Claude (Opus 4.7, Anthropic) were used to assist with editing, refactoring, and debugging portions of the analysis code. All AI-assisted code was reviewed, edited as needed, executed, and validated by the authors before use in the reported analyses.

#### Lead Contact

Requests for further information and resources should be directed to and will be fulfilled by the lead contact

#### Materials Availability

This study did not generate new unique reagents.

### Data and Code Availability

- CITE-seq data generated in this study, including processed gene-expression and antibody-derived tag count matrices, cell-level metadata, and processed single-cell Seurat objects, will be deposited in the NCBI Gene Expression Omnibus (GEO). Raw sequencing data will be deposited in the associated Sequence Read Archive (SRA). Accession numbers will be provided before publication.
- All original code used for the analyses in this study will be deposited at Zenodo (https://www.zenodo.org/) and made publicly available with a unique digital object identifier (DOI) before publication. The code will also be made available through GitHub (https://www.github.com/).
- Any additional information required to reanalyze the data reported in this paper is available from the lead contact upon request.

### Declaration of generative AI and AI-assisted technologies in the manuscript preparation process

During the preparation of this work, the authors used ChatGPT (OpenAI) and Claude (Anthropic) to assist with editing and refinement selected manuscript text. After using these tools, the authors reviewed and edited the content as needed and take full responsibility for the content of the published article.

