## Supplemental Figures for "Intravenous BCG Elicits a Sustained Myeloid IFNG Response Signature Associated with Early TB Control"

### Supplemental Figure 1

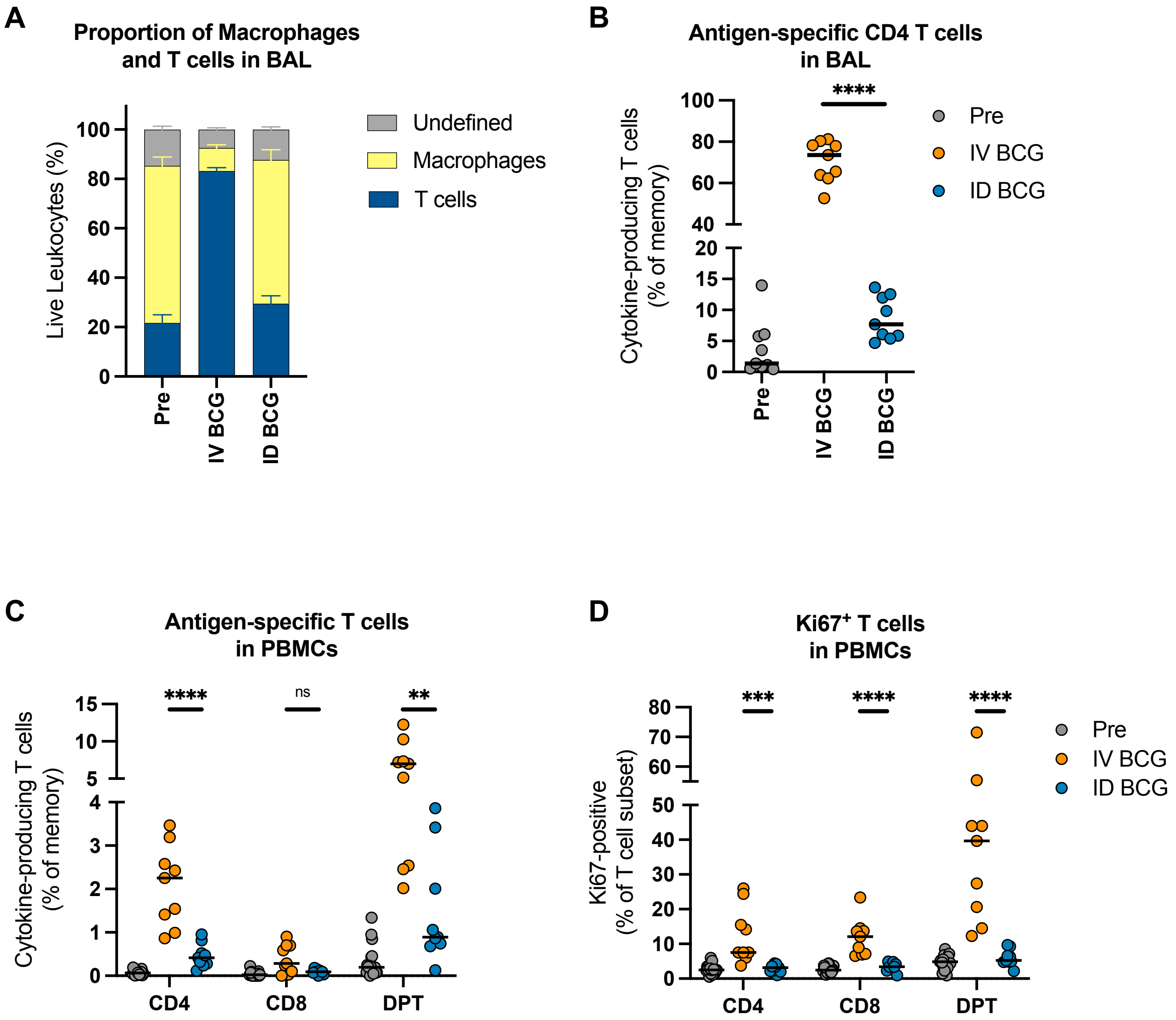

**Supplemental Figure 1. Cellular composition and CD4 T cell responses in BAL and PBMCs after BCG vaccination**

(**A**) Flow cytometry was used to analyze the relative proportion of macrophages and T cells (percent of live CD45+ leukocytes) in BAL samples collected before (Pre) and after (8 weeks) IV or ID BCG vaccination. Bars show the mean proportions of 9 animals per group (with SEM).

(**B**) Mtb whole cell lysate (WCL)-specific T cell responses in BAL after vaccination, measured by intracellular cytokine staining. Shown are the percentages of CD4 T cells in BAL producing any combination of IFNG, IL2, TNFA, and IL17A before (Pre) and after (8 weeks) IV or ID BCG vaccination. Symbols represent individual macaque responses (media alone background subtracted) with medians. ****$P$ <0.0001 (Mann-Whitney).

(**C**) Mtb whole cell lysate (WCL)-specific T cell responses in PBMCs after vaccination, measured by intracellular cytokine staining. Shown are the percentages of memory CD4+CD8-, CD8+, and CD4+CD8+ double positive (DP) T cells PBMCs producing any combination of IFNG, IL2, TNFA, and IL17A before (Pre) and after (4 weeks) IV or ID BCG vaccination. Symbols represent individual macaque responses (media alone background subtracted) with medians. ****$P$ <0.0001; **$P$ = 0.0012; ns, not significant (Mann-Whitney).

(**D**) The percent of T cells in PBMCs expressing Ki67, measured by flow cytometry. T cell subsets shown include CD4+CD8-, CD8+, and CD4+CD8+ double positive (DP) T cells. Symbols represent individual macaque responses with medians. ****$P$ <0.0001; ***$P$ = 0.002 (Mann-Whitney).

### Supplemental Figure 2

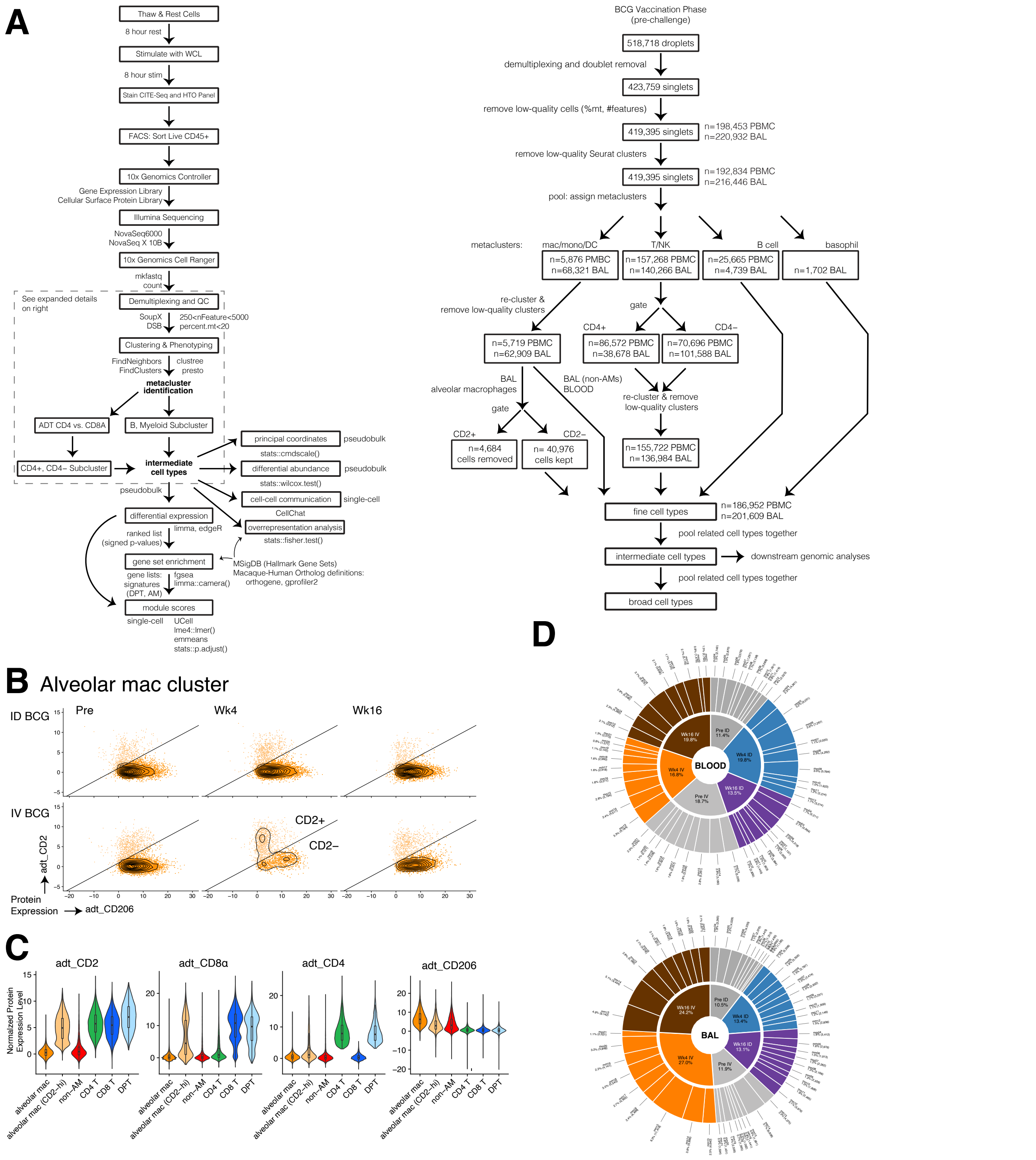

**Supplemental Figure 2. Detailed overview of experimental and bioinformatic pipeline and select BAL alveolar macrophage cell surface expression markers**

(**A**) Flow chart outlining thawing, staining, and sorting protocol prior to running 10x Genomics Controller for GEM bead generation. Flow chart continues with sequencing specifications, 10x CellRanger demultiplexing and brief clustering details followed by analysis approaches and respective R packages used (left). Expanded detailed summary of demultiplexing and quality control steps followed by iterative clustering to annotate multiple cell clustering resolutions (right).

(**B**) Cell surface protein expression (ADT) of CD2 vs CD206 on BAL alveolar macrophages, split by vaccination group and timepoint, with line used to assign cells into CD2+ or CD2– alveolar macrophage subclusters. Only CD2– alveolar macrophages were retained for downstream analysis (bottom).

(**C**) Cell surface protein expression (ADT) of CD2, CD8α, CD4, and CD206 for select alveolar mac and T cell type (meta) clusters.

(**D**) Proportion of total analyzed cells from each of the 18 macaques at each of the 3 timepoints, grouped by vaccine and timepoint, for blood (top) and BAL (bottom).

### Supplemental Figure 3

#### A Cell type cluster markers - resolution: metacluster

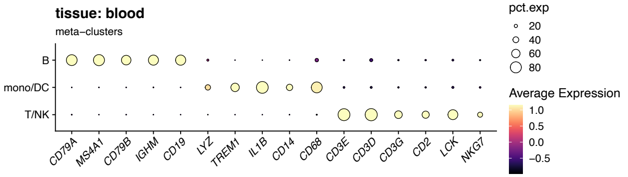


#### B Cell type cluster markers - resolution: broad

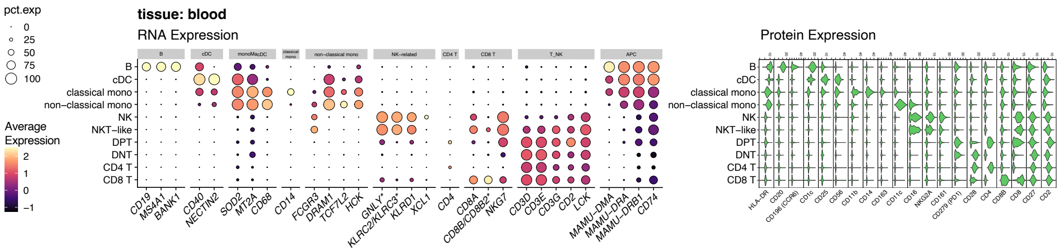


#### C Cell type cluster markers - resolution: intermediate

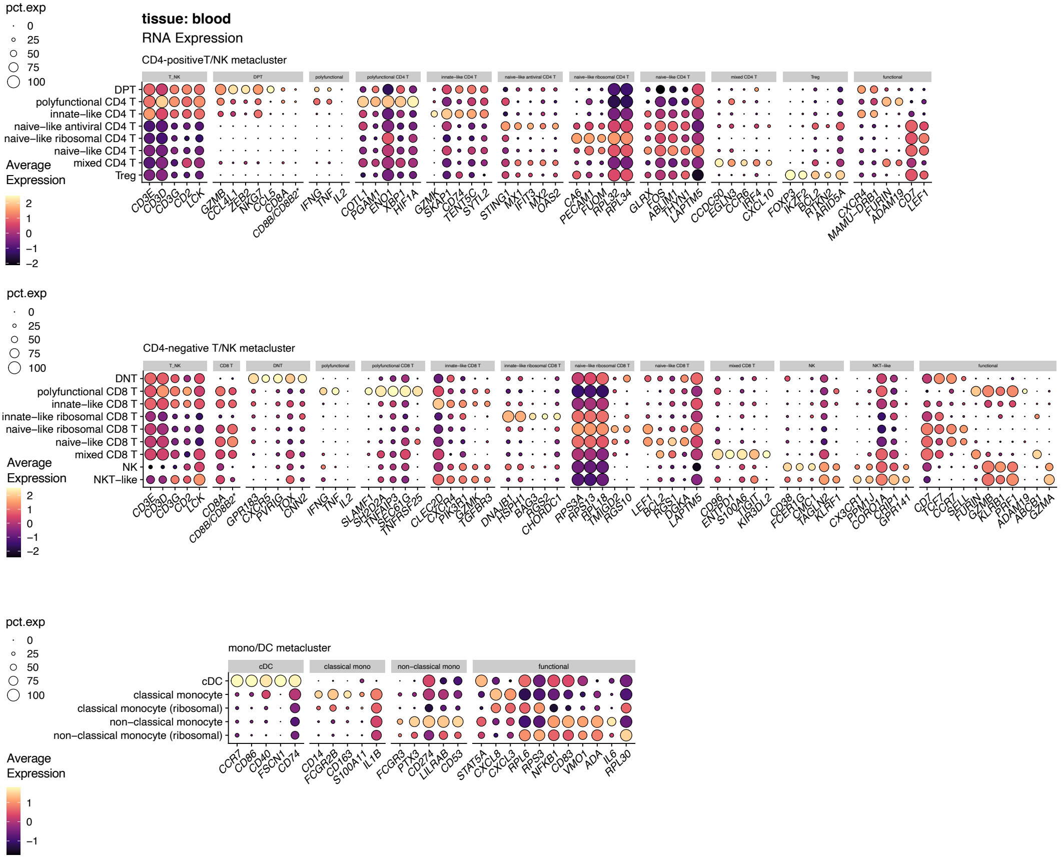

**Supplemental Figure 3. Curated marker expression for PBMC cell type annotation during the vaccination phase**

(**A**) Expression of select marker genes used to distinguish and annotate metaclusters in the vaccination phase PBMCs: B, mono/DC, and T/NK cells.

(**B**) RNA expression (left) and cell surface protein expression (ADT, right) of selected markers used to distinguish and annotate broad resolution vaccination phase PBMC cell types.

(**C**) RNA expression of selected markers used to distinguish and annotate intermediate resolution vaccination phase PBMC cell types among the CD4-positive T/NK metacluster (top), CD4-negative T/NK metacluster (middle), and mono/DC metacluster (bottom).

### Supplemental Figure 4

#### A IV-specific Wk4 vs Pre DPT signature genes
#### Top 100 (by significance)

| | | | |
|---|---|---|---|
| *ABCC1* | *HIVEP3* | *PGAM1* | *SEC61B* |
| *ADAM19* | *HM13* | *PGAM4** | *SEC61G* |
| *ADTRP* | *HNRNPAB* | *PHLDA1* | *SERBP1* |
| *ARF6* | *HSPA9* | *PICALM* | *SFPQ* |
| *ARPC5L* | *IL21R* | *PIGT* | *SFT2D1* |
| *BATF* | *ILF2* | *PIM3* | *SLAMF1* |
| *BCL2L1* | *LAPTM4B* | *PITRM1* | *SLC1A5* |
| *BHLHE40* | *LCP2* | *PKIG* | *SLC9A7* |
| *C1QBP* | *LDHA* | *PLAGL2* | *SMS* |
| *CALM1** | *LDHA** | *POMP* | *SND1* |
| *CBLB* | *LRRFIP1* | *PPM1M* | *SPAG9* |
| *CD2* | *MAP2K3* | *PRDX1* | *SRPRB* |
| *CD3D* | *MAST4* | *PRDX6* | *SRSF2* |
| *CD53* | *MAT2A* | *PTP4A2* | *SURF4* |
| *CD109* | *MLLT6* | *RAB11FIP1* | *TNFRSF8* |
| *CD274* | *MRPL9* | *RANBP1* | *TNFRSF25* |
| *CDK6* | *MSC* | *RAP1A* | *TRAF3* |
| *COTL1* | *MTHFD2* | *RBM17* | *TRIB1* |
| *CTSB* | *MYBBP1A* | *RBPJ* | *TRIM72* |
| *DAD1* | *NDFIP1* | *REV3L* | *U2AF1** |
| *DBNDD2* | *ODC1* | *RGCC* | *UFM1* |
| *EED* | *PARK7* | *RILPL2* | *VSIG8* |
| *EIF4A1* | *PARP6* | *RNF19A* | *XBP1* |
| *EIF4G1* | *PFKFB3* | *RORC* | *ZNRF1* |
| *ETV7* | *PFKP* | *RUNX1* | ENSMMUG00000061824 |

#### C DPT signature genes: Overrepresentation Analysis (ORA)

padj.BH < 0.10; pathway overlap > 5

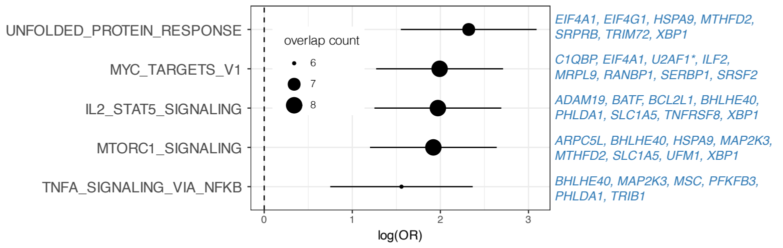


## D

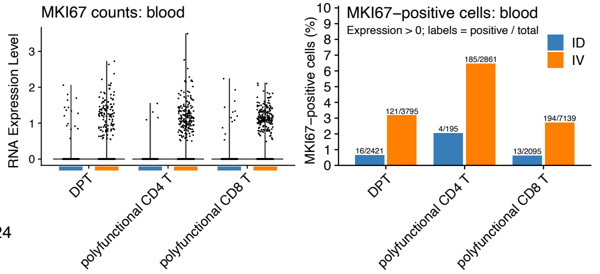


## B

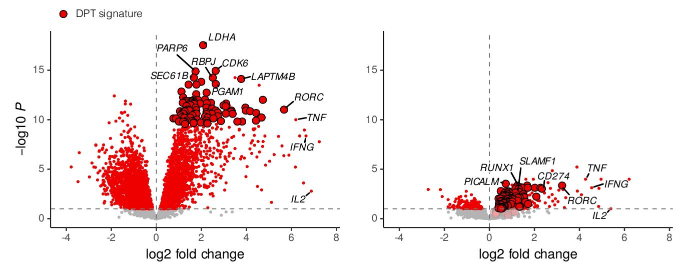


| signature | DE genes, n/N (% of signature) Wk4 vs. Pre | Wk16 vs. Pre |
|---|---|---|
| DPT100 | 100/100 (100%) | 79/100 (79.0%) |
| T1T17 | 39/45 (86.7%) | 20/45 (44.4%) |
| module2_hi | 42/47 (89.4%) | 26/47 (55.3%) |

## E

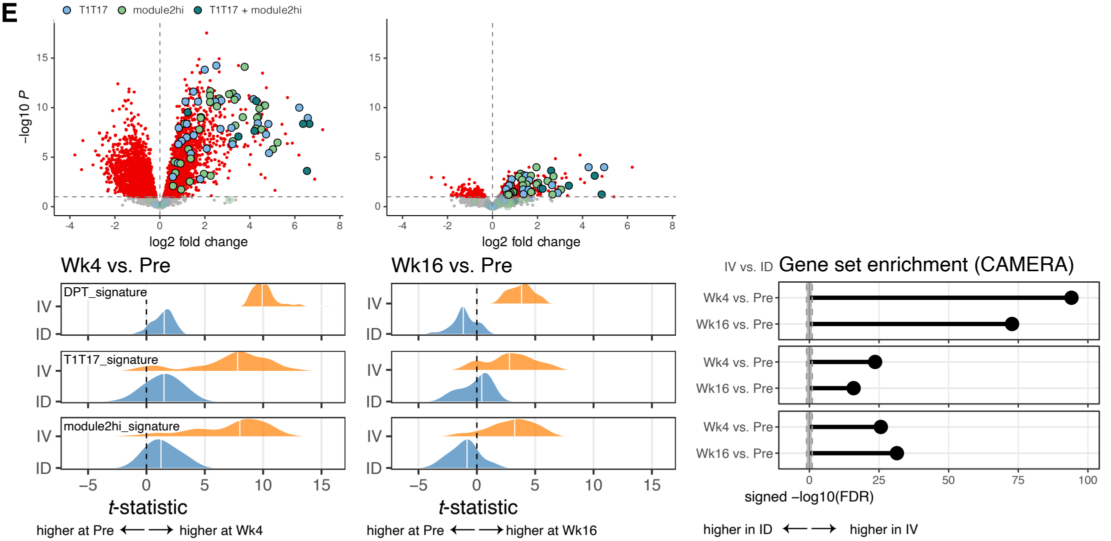

**Supplemental Figure 4. Transcriptional signature of IV BCG responsive CD4+CD8α+ double positive T cells from vaccination phase blood**

(**A**) List of the top 100 genes (ranked by –log10(BH-adjusted *P* value) upregulated in DPT cells (Wk4 vs. Pre) following IV BCG (BH FDR $\leq$ 0.1; logFC > 0.25), excluding genes also upregulated following ID BCG at a relaxed threshold (BH FDR $\leq$ 0.4 logFC > 0.25) comprising an "IV BCG-specific DPT signature".

(**B**) Volcano plot of differential gene expression (red; BH FDR $\leq$ 0.1) in DPT cells from IV BCG samples at Wk4 vs Pre (left) and Wk16 vs Pre (right), with the IV-specific DPT signature genes outlined in black and *IFNG*, *TNF*, and *IL2* also labeled. As indicated in the table to the right, 79 of the 100 DPT signature genes remained significantly upregulated at Wk16 (vs. Pre).

(**C**) Overrepresentation analysis (ORA) of Hallmark pathways among the DPT signature genes (one-sided Fisher's exact test; BH FDR < 0.1; Hallmark gene sets shown only when >5 signatures genes overlapped), with overlapping pathway genes listed on the right.

(**D**) Ki67 gene expression within DPT, polyfunctional CD4 and polyfunctional CD8 T cells at Wk4 after IV or ID BCG vaccination in PBMCs. Shown are Ki67 RNA expression level (left) and percent Ki67-positive cells (right) for each vaccine group.

(**E**) Volcano plots show differential gene expression in DPT cells (red) from IV BCG samples as in (**B**). Genes shared with the top 50 most significant DE genes in previously published Th1/Th17 gene modules ("module 2hi" and "T1T17")[10,23,27] are indicated by color (blue/green) and outline and shown for Wk4 and Wk16. Number of shared genes among annotated genes in the DPT dataset are shown for each module in table to the right. Ridgeline plots show *t* statistics for DPT signature (top), T1T17 signature (middle), and module2hi signature (bottom) in DPT cells from IV and ID samples at Wk4 (vs. Pre) and Wk16 (vs. Pre). Solid white lines denote each group median. Statistical analysis (right, lollipops): CAMERA competitive gene-set testing of each signature at Wk4 and Wk16, with significance given by $|\log_{10}FDR| > 1$.

### Supplemental Figure 5

**A Vaccine Phase GSEA: BLOOD intermediate resolution cell types**

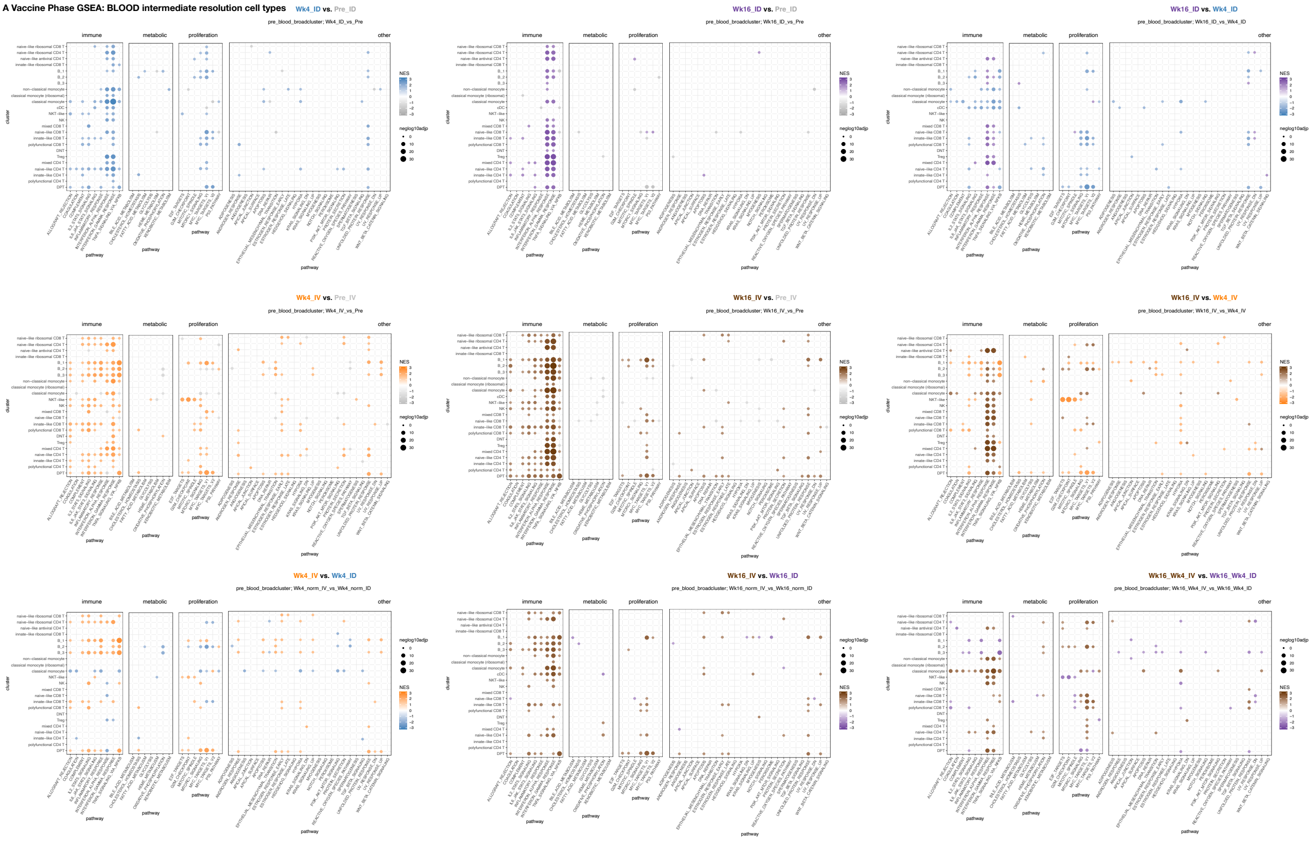

**Supplemental Figure 5. Comprehensive Hallmark pathway gene set enrichment analysis of vaccination phase PBMCs**

(**A**) Dot plots depicting gene set enrichment analyses of all Hallmark pathways from blood vaccination phase intermediate resolution cell types. The top row corresponds to: ID specific contrasts for Wk4 vs Pre, Wk16 vs Pre, and Wk16 vs Wk4; the middle row corresponds to IV specific contrasts for Wk4 vs Pre, Wk16 vs Pre, and Wk16 vs Wk4; and the bottom row corresponds to difference-of-differences contrasts for IV vs. ID: Wk4 vs Pre, Wk16 vs Pre, and Wk16 vs Wk4.

### Supplemental Figure 6

#### A Cell type cluster markers - resolution: metacluster

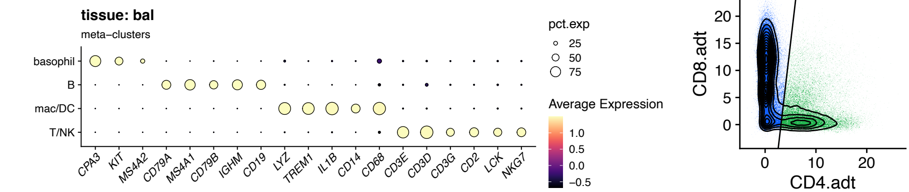


#### B Cell type cluster markers - resolution: broad

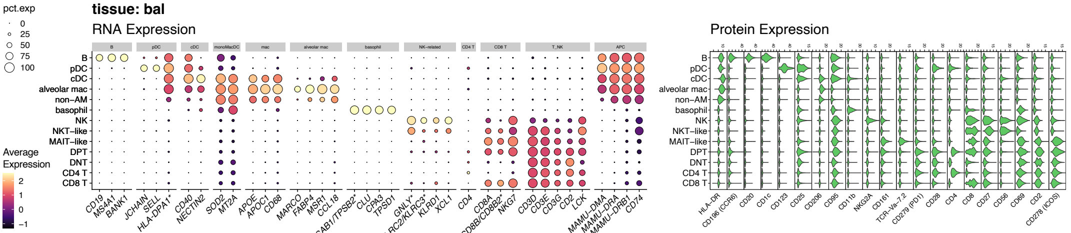


#### C Cell type cluster markers - resolution: intermediate

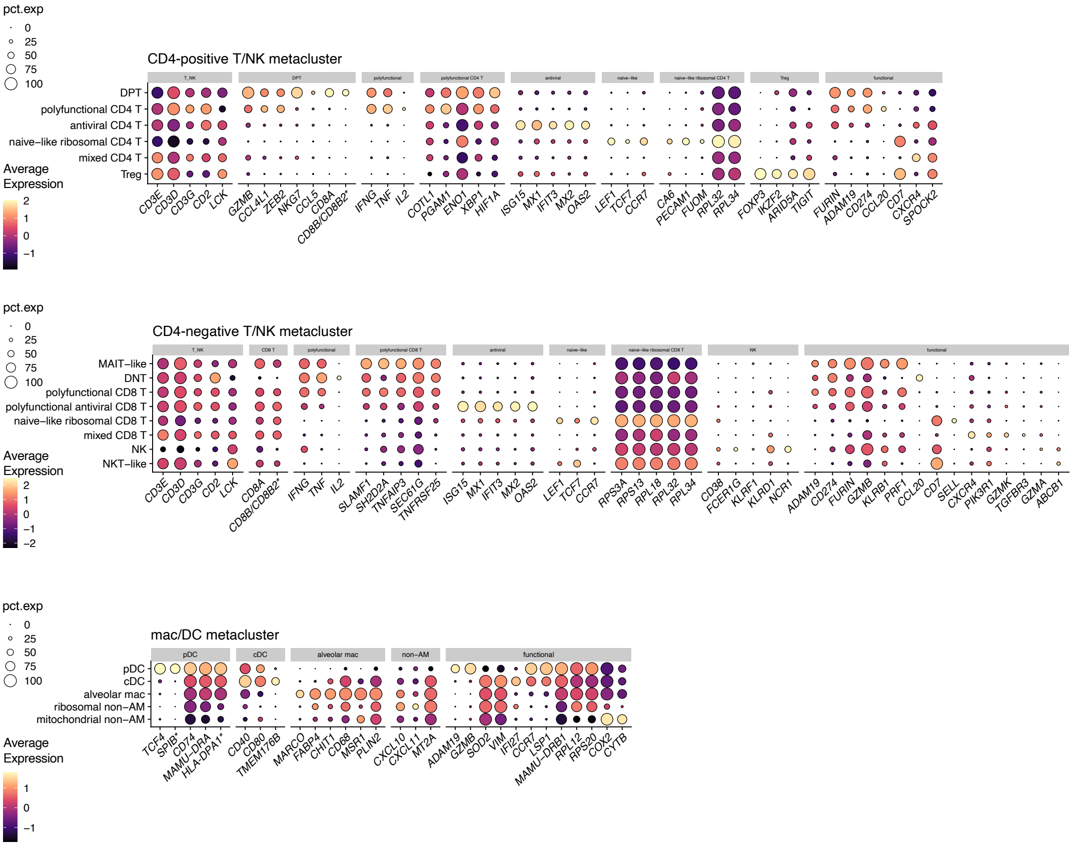

**Supplemental Figure 6. Curated marker expression for BAL cell type annotation during the vaccination phase**

(**A**) Expression of select marker genes used to distinguish and annotate metaclusters: basophil, B, mac/DC, and T/NK in the vaccination phase BAL (left). Scatterplot of cell surface protein expression of CD8α vs. CD4 with line used to assign downstream T/NK subclusters (right).

(**B**) RNA expression (left) and cell surface protein expression (right) of selected markers used to distinguish and annotate broad resolution vaccination phase BAL cell types.

(**C**) RNA expression of selected markers used to distinguish and annotate intermediate resolution vaccination phase BAL cell types among the CD4-positive T/NK metacluster (top), CD4-negative metacluster, and macrophage/DC metacluster.

### Supplemental Figure 7

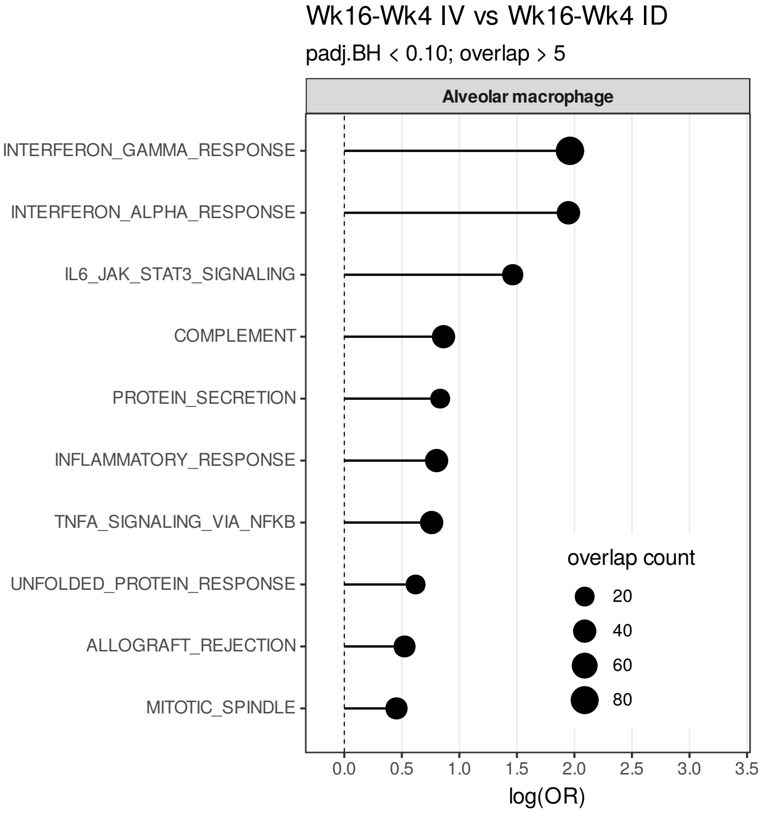

**Supplemental Figure 7. Hallmark gene set overrepresentation analysis of the IV vs. ID response in BAL alveolar macrophages**

Hallmark gene sets overrepresented among upregulated DEGs (BH FDR < 0.1; logFC > 0.25) for the IV vs. ID, Wk16 vs. Wk4 contrast in BAL AMs. One-sided Fisher's exact test; BH FDR < 0.1; Hallmark gene sets shown only when > 5 DEGs overlapped.

### Supplemental Figure 8

**A Vaccination Phase GSEA: BAL intermediate resolution cell types**

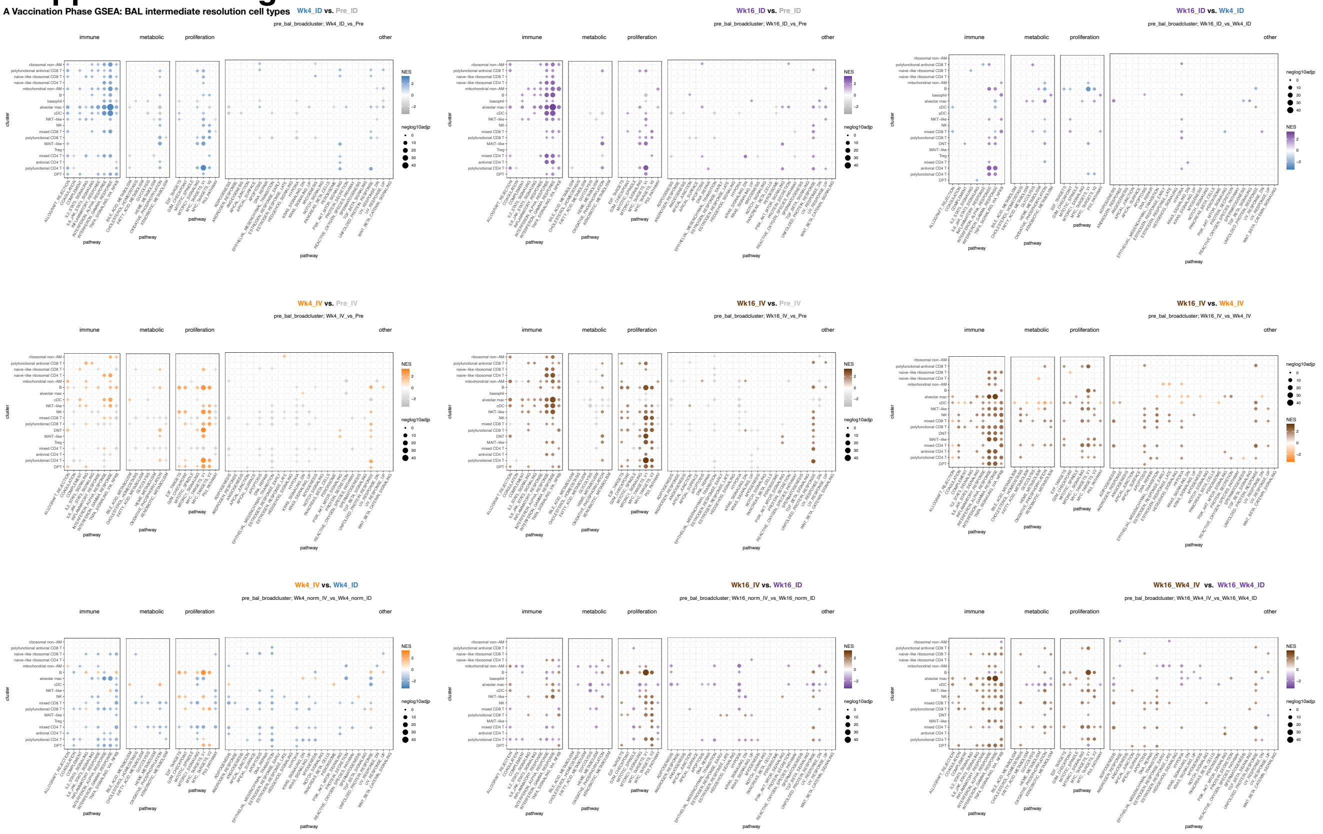

**Supplemental Figure 8. Comprehensive Hallmark pathway gene set enrichment analysis of vaccination phase BAL cells**

(**A**) Dot plots depicting gene set enrichment analyses of all Hallmark gene sets from BAL vaccination phase intermediate resolution cell types. The top row corresponds to: ID specific contrasts for Wk4 vs Pre, Wk16 vs Pre, and Wk16 vs Wk4; the middle row corresponds to IV specific contrasts for Wk4 vs Pre, Wk16 vs Pre, and Wk16 vs Wk4; and the bottom row corresponds to difference-of-differences contrasts for IV vs. ID: Wk4 vs Pre, Wk16 vs Pre, and Wk16 vs Wk4.

### Supplemental Figure 9

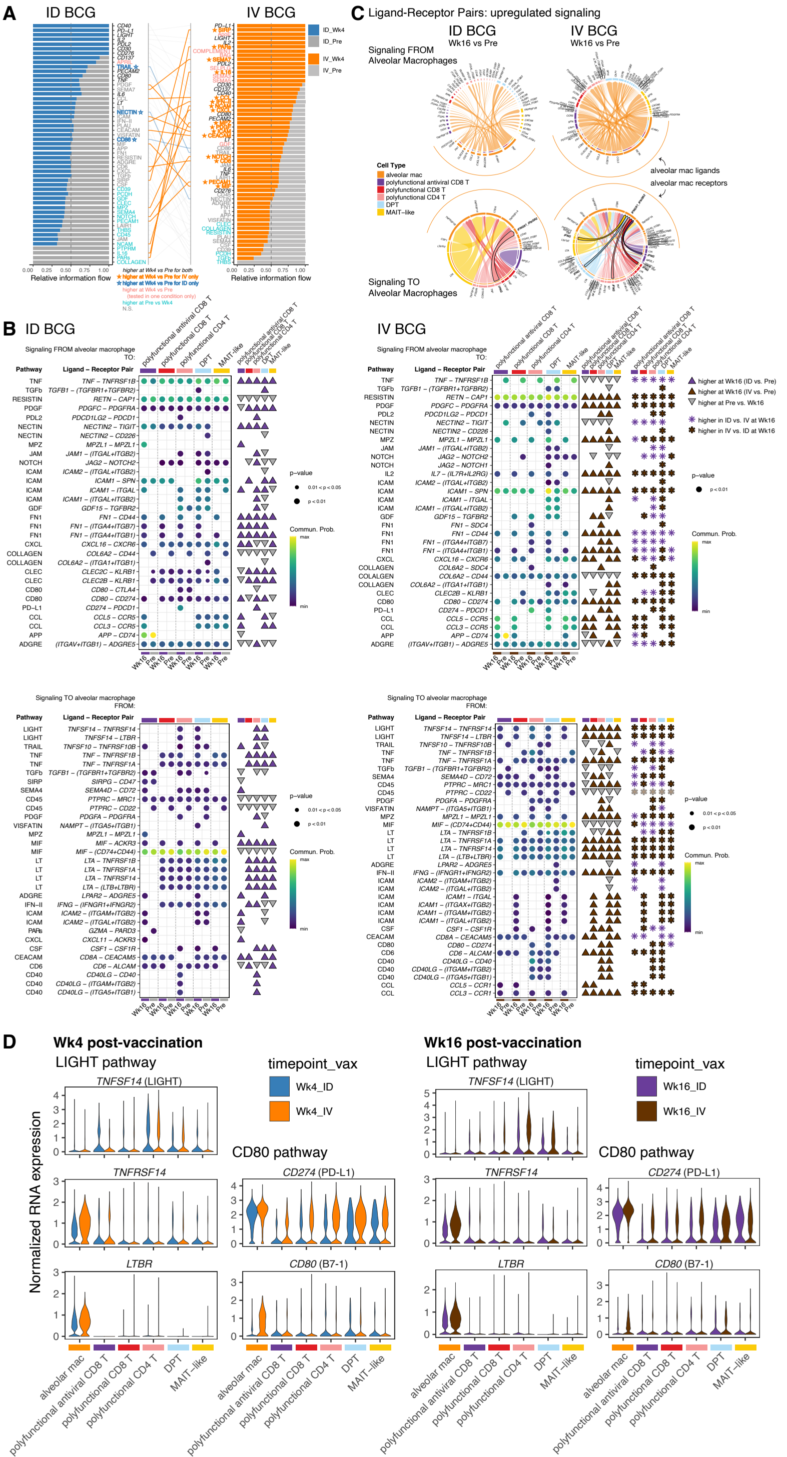

**Supplemental Figure 9. Expanded cell-cell, inferred ligand-receptor communication signaling in the BAL.**

(**A**) Bar graph showing the rank-ordered relative information flow (Wk4 vs Pre) of each assessed cell communication pathway for ID BCG (left) and IV BCG (right) across all cell types. Stars denote signaling pathways with significantly higher relative information flow at Wk4 vs. Pre (CellChat rankNet test, $P < 0.05$), with blue for ID and orange for IV. Colored lines link the same pathway across groups, and pathway names are colored in the group with higher relative induction of information flow. The greater number of orange lines suggest IV vaccination induces enhanced cellular communication from Wk4 to Pre compared to ID. Signaling pathways in turquoise were higher at Pre within their respective BCG vaccination group. Black italicized signaling pathways were higher at Wk4 for both groups, while those greyed out were not significant in either group.

(**B**) CellChat-inferred ligand-receptor communication probabilities between BAL alveolar macrophages and selected T cell subsets at Wk16 and Pre in ID BCG (left) and IV BCG (right). Top, signaling from alveolar macrophages to T cell subsets; bottom, signaling from T cell subsets to alveolar macrophages. Colored circles represent significant inferred ligand-receptor interactions (CellChat permutation test, $P < 0.05$); circle color indicates communication probability and size indicates $P$ value. Colored triangles indicate the direction of differences between Wk16 and Pre within each vaccine group identified in separate CellChat comparisons. Colored asterisks indicate differences between vaccination groups at Wk16. Faded asterisks denote Wk16 between-group differences in which the higher-probability group decreased from Pre to Wk16.

(**C**) Chord diagrams show CellChat-inferred AM–T cell interactions associated with differential ligand expression between Pre and Wk16 (Wilcoxon rank-sum test, $P < 0.05$; $|\text{ligand logFC}| > 0.05$) for ID and IV BCG. Chords connect ligand and receptor genes to their corresponding source and target cell populations, and chord width represents relative inferred communication strength within each chord diagram. IFN-II pathway ligand-receptor pairs (T cells to AMs, bottom) are bolded for emphasis.

(**D**) Violin plots of ligand-receptor gene expression in BAL alveolar macrophages, polyfunctional antiviral CD8 T, polyfunctional CD8 T, polyfunctional CD4 T, DPT, and MAIT-like cells for the LIGHT pathway (TNFSF14, TNFRSF14, LTBR) and the CD80 pathway (PDL1, CD80) at Wk4 (left) in ID (blue) or IV (orange). The corresponding panels for Wk16 are on the right for ID (purple) and IV (brown).

### Supplemental Figure 10

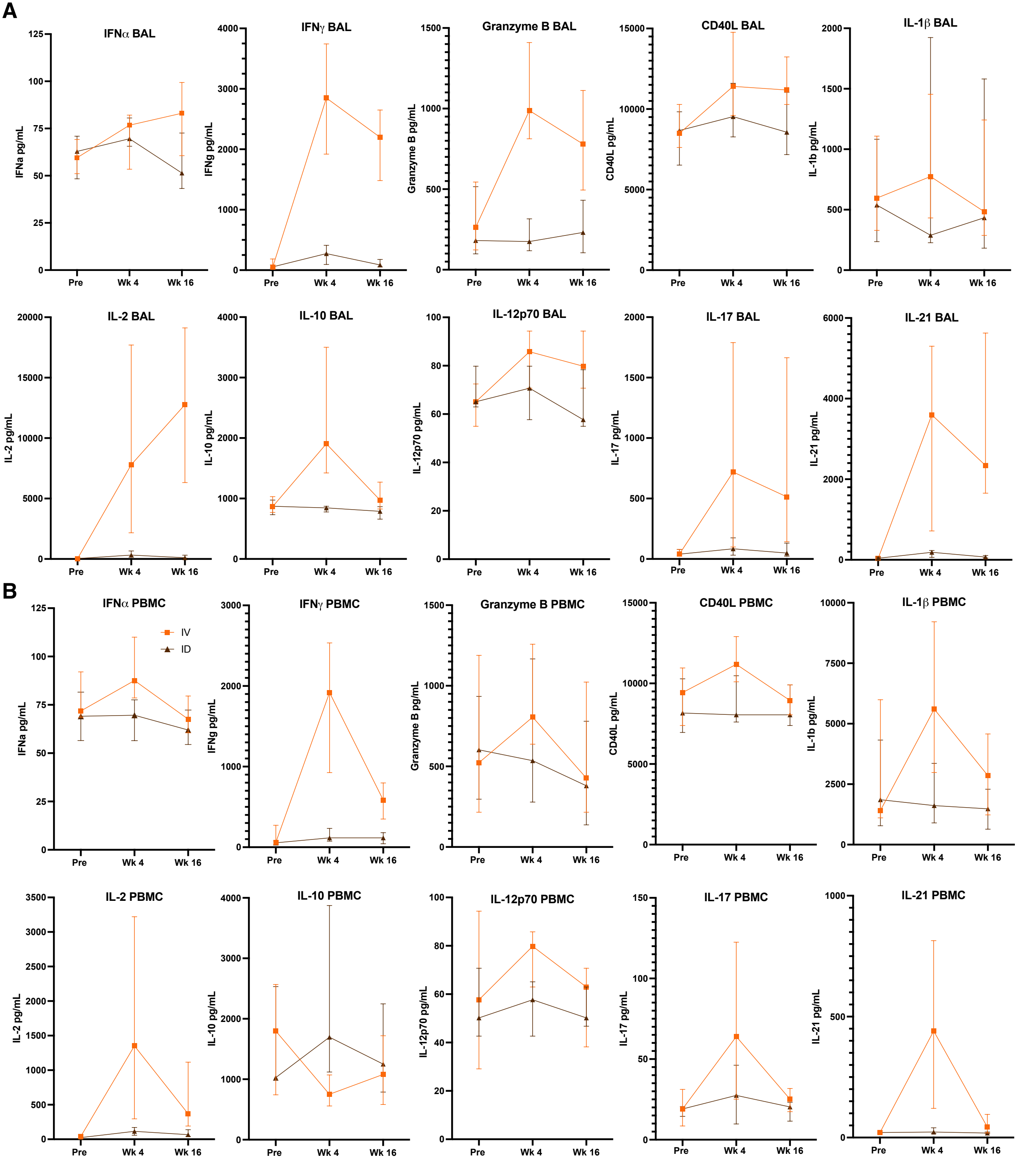

**Supplemental 10. Cytokine production after *in vitro* stimulation with Mtb WCL**

Cytokines measured from cell culture supernatants collected from BAL (**A**) and PBMC (**B**) cells that were stimulated for 8hrs with Mtb WCL for subsequent single cell RNA processing; a custom BioTechne NHP XL Cytokine Performance Kit was used. Shown are the median values of 9 macaques per group per timepoint with 95% confidence intervals.

#### Supplemental Figure 11

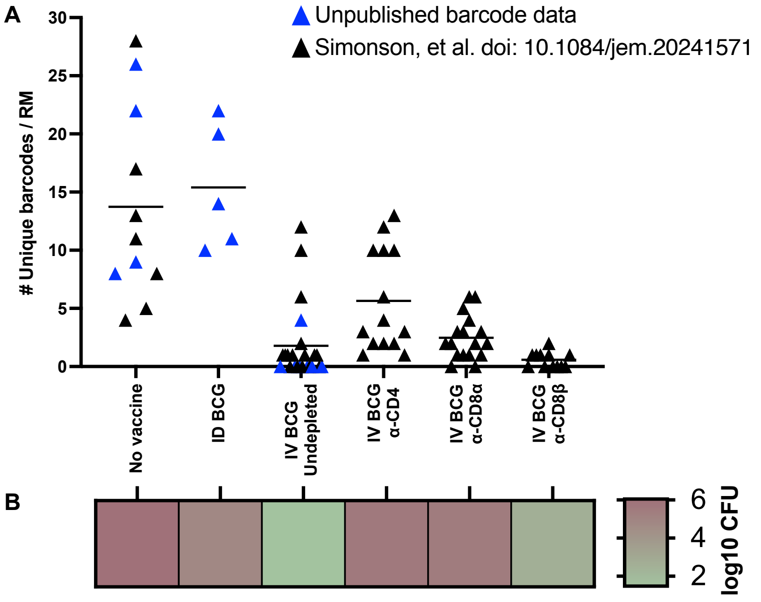


Summary of CFU data from
Darrah, et al. doi: 10.1038/s41586-019-1817-8
Simonson, et al. doi: 10.1084/jem.20241571

**Supplemental 11. Quantification of Mtb establishment after infection of IV or ID BCG immunized macaques**

In animals that have been challenged with a barcoded strain of Mtb Erdman, the number of individual bacilli that establish TB infection can be determined by the quantifying the number of unique barcodes from Mtb CFUs recovered from all tissues at study endpoint (8–12 weeks post-challenge). Panel (**A**) shows the total number of unique barcodes identified per RM and combines data from two prior studies: black symbols represent barcode data published previously[13]; blue symbols represent unpublished barcode data from a previously published efficacy study in macaques[10] from which a limited number of samples per group were analyzed (see Table S2 for animal information). In the ID BCG group, n=3 animals were immunized with the standard dose (~$5\times10^5$ CFU), and n=2 animals received a high dose of ID BCG (~$5\times10^7$ CFU) that was matched to the IV group. Some groups of IV BCG immunized animals received antibodies to deplete CD4 T cells, or CD8α- or CD8β-expressing cells 5 months after vaccination and one month before (and throughout) Mtb challenge, as described[13]. Values of zero represent animals with no detectable CFU at endpoint, bars are medians. (**B**) Heat map depicting the median CFU (log10) at endpoint for all animals in each vaccine group from both published studies[10,13].

### Supplemental Figure 12

A

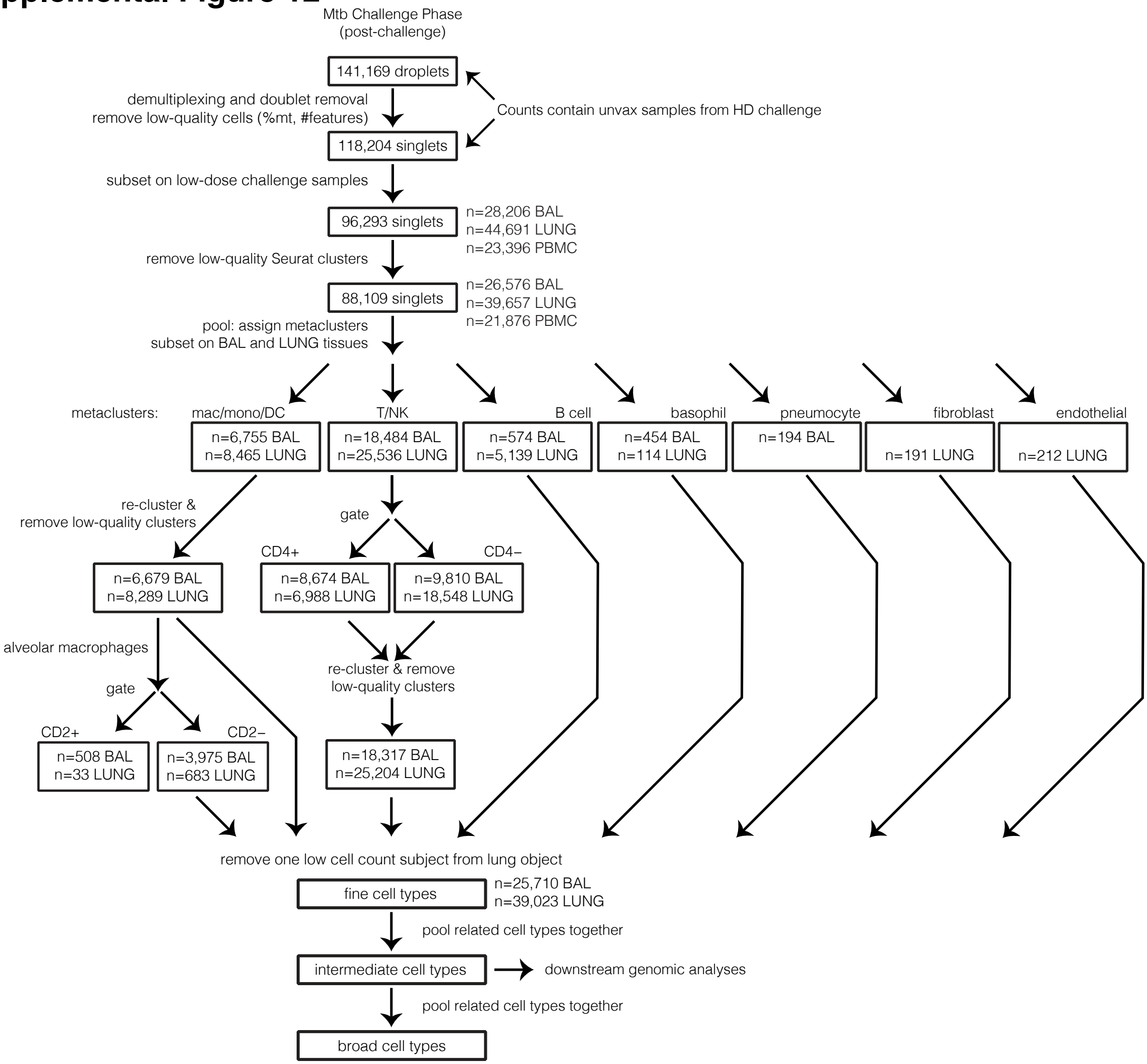


B

Challenge phase

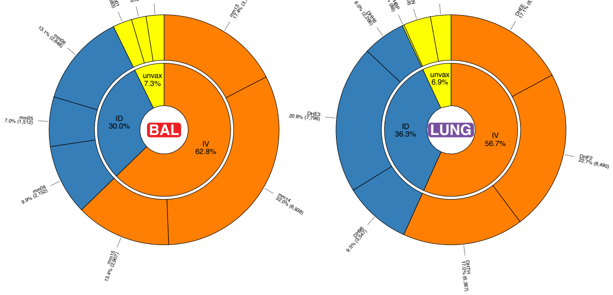

**Supplemental Figure 12. Overview of quality control and clustering iterations for post-challenge single cell objects**

(**A**) Flow chart depicting cell counts at each stage of quality control for challenge phase single cell samples. Single-cell barcodes were demultiplexed and doublets removed via hashtag expression followed by the removal of low-quality cells as determined by percentage of mitochondrial reads and number of unique genes expressed (features). After initial clustering, poor quality clusters were removed prior to re-clustering and cellular annotation. Number of cells included in each metacluster for BAL and lung single cell objects are denoted, with the downstream workflow disclosing the iterative clustering process.

(**B**) Proportion of total analyzed cells from each of the 9 macaques challenged with Mtb, grouped by vaccine, for BAL and lung in the post-challenge phase (D11).

### Supplemental Figure 13

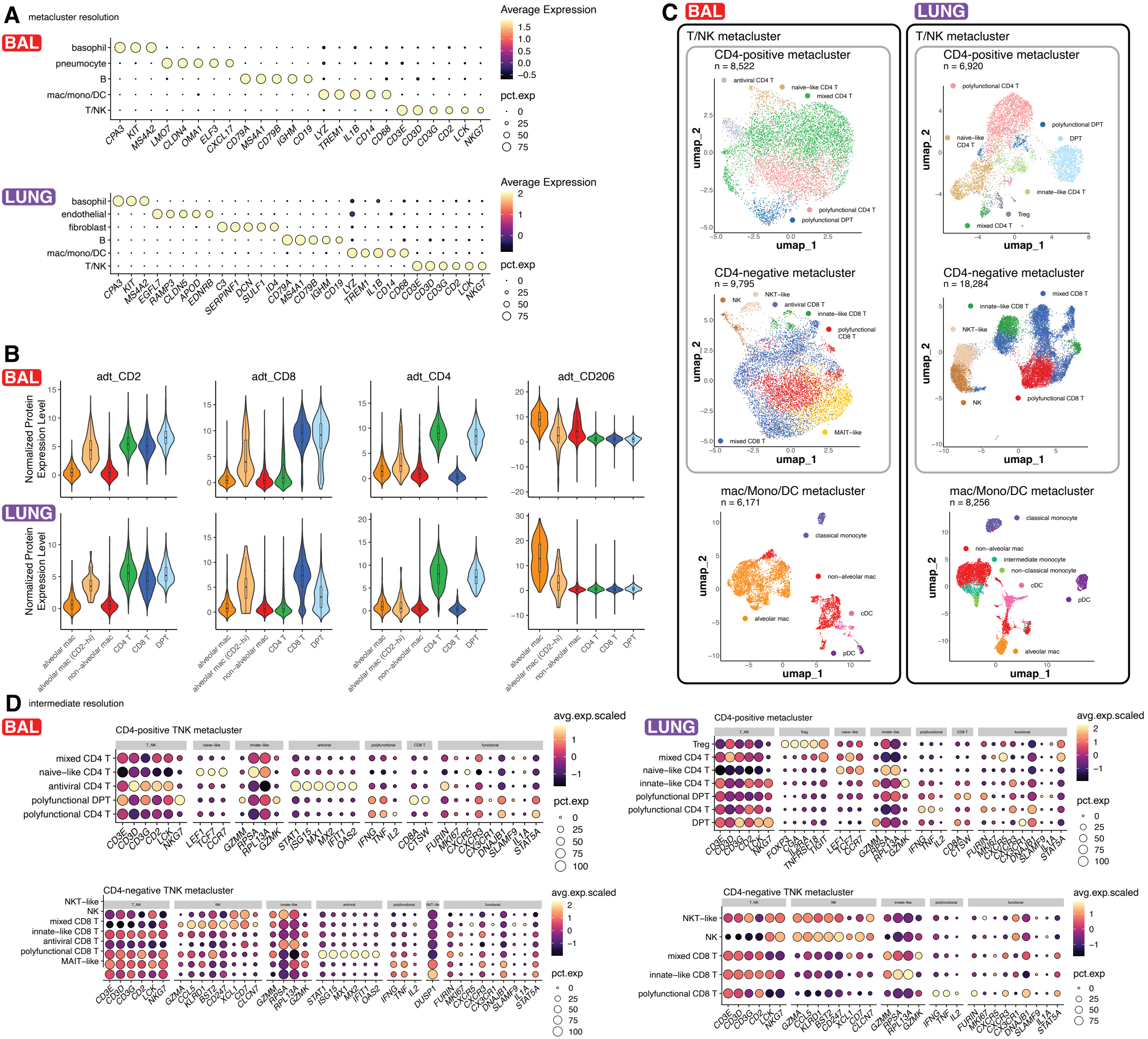

**Supplemental Figure 13. Curated RNA marker expression, select cell surface marker expression, and UMAPs for challenge phase BAL and lung cells.**

(**A**) Curated gene markers used to split post-challenge BAL (top) and lung (bottom) metaclusters. Post-challenge BAL metaclusters include basophil, pneumocytes, B, mac/mono/DC, and T/NK clusters while lung cells contain basophil, endothelial, fibroblast, B, mac/mono/DC, and T/NK cell clusters. Endothelial cells and fibroblasts were detected despite CD45+ flow sorting.

(**B**) Violin plots for cell surface expression in post-challenge BAL (left) and lung (right) for markers in descending column order: CD2, CD3, CD4, CD8α, and CD206.

(**C**) UMAPs for BAL (left) and lung (right) intermediate resolution cell type clusters in the CD4-positive metacluster (top), CD4-negative metacluster (middle), and mac/mono/DC metacluster (bottom).

(**D**) Dot plots for curated gene expression markers used to distinguish and annotate intermediate cell types within CD4-positive, CD4-negative, and mac/mono/DC metaclusters in BAL and lung tissues.

### Supplemental Figure 14

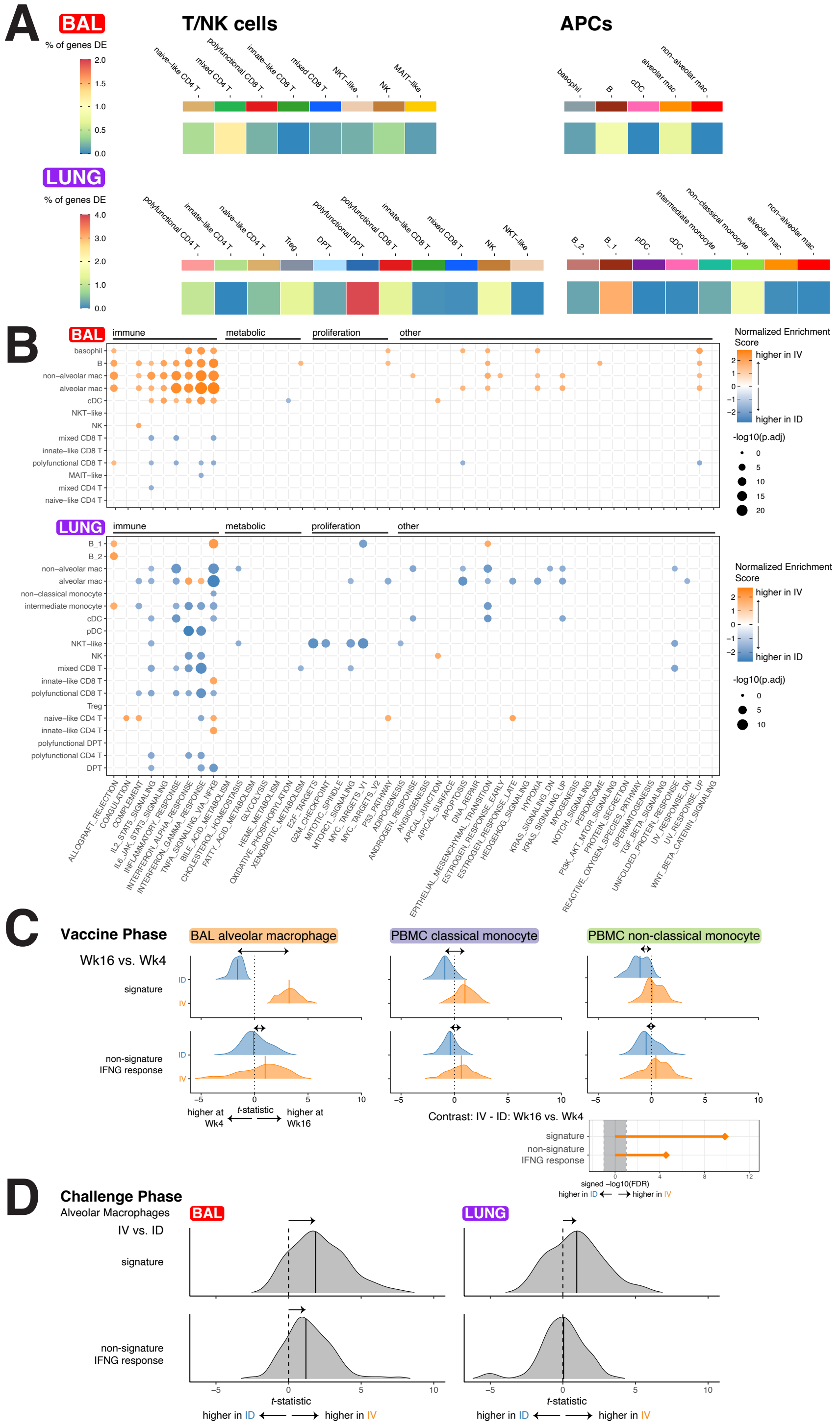

**Supplemental Figure 14. Differential gene expression, gene set enrichment analysis, t statistic linear modeling outputs, and CAMERA enrichment analysis for post-challenge BAL and lung.**

(**A**) Heatmap for BAL (left) and lung (right) depicting the percent of differentially expressed genes (BH FDR < 0.1) between IV vs. ID within APCs (top) or T cell subsets (bottom). Analysis was limited to cell types which met cell thresholding parameters in both vaccination groups.

(**B**) Dot plots depicting gene set enrichment analyses of all Hallmark gene sets from post-challenge BAL (top) and lung (bottom) intermediate resolution cell types.

(**C**) Ridgeline plots of moderated *t* statistics for the 61-gene IFNG response signature and non-signature IFNG response gene sets in vaccination phase BAL alveolar macrophages (left), PBMC classical monocytes (middle), and PBMC non-classical monocytes (right) for the Wk16 vs Wk4 contrast. Arrows depict the difference in the *t* statistic medians for ID vs. IV. Statistical analysis (bottom right), CAMERA competitive gene-set testing (lollipops) of the 61-gene IFNG response signature enrichment in blood non-classical monocytes (IV vs. ID; Wk16 vs Wk4) with significance threshold of |log10FDR| >1.

(**D**) Ridgeline plots of moderated *t* statistics from pseudobulk DE analysis for the 61-gene IFNG response signature (top) and IFNG response non-signature (bottom) gene sets in post-challenge BAL AMs (left) or lung AMs (right) for the IV vs. ID contrast. Increased signature enrichment in IV is depicted by a rightward shift of the median line from the dashed center line.

#### Supplemental Figure 15

A Wk4: Effect of Stimulation on Cytokine Production

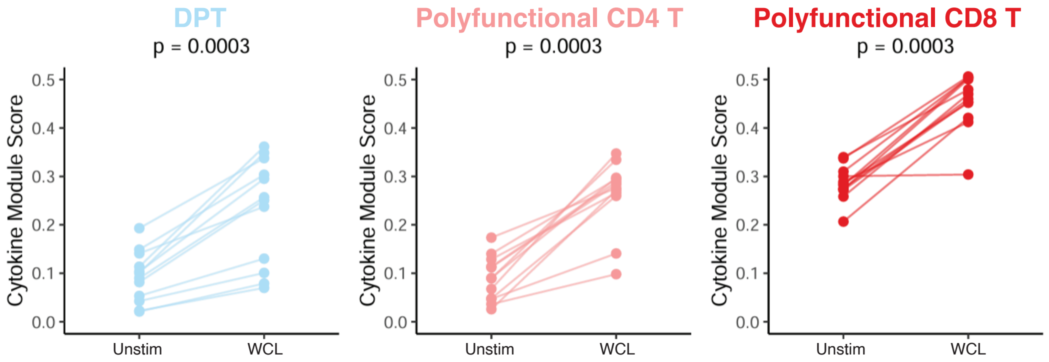


*Cytokine Module = IFNg, TNF, IL2, IL17a, GZMB, PRF1*

B Discovery Cohort DPT Week 4 Top100 Signature

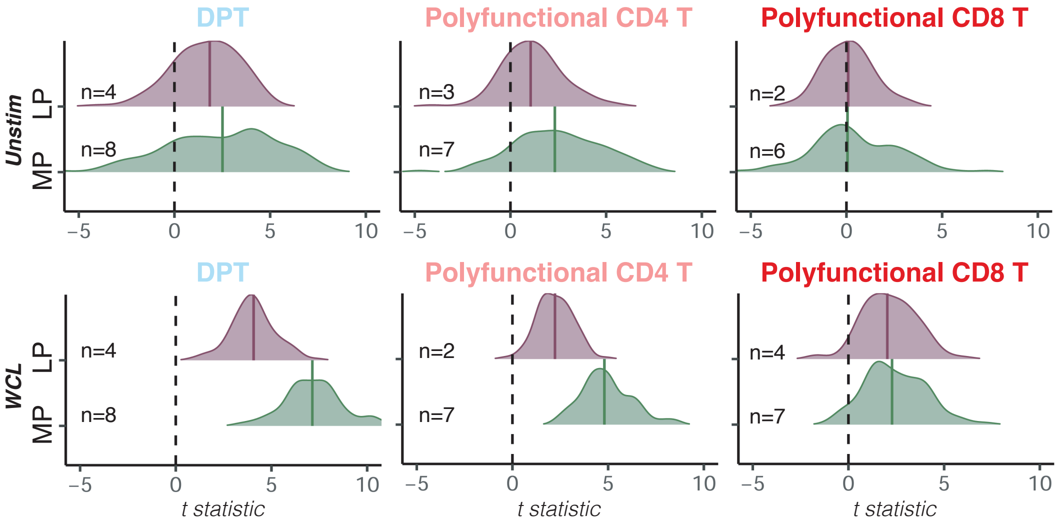

**Supplemental Figure 15. Effect of antigen stimulation on the DPT signature across T cell subsets in an orthogonal dataset.**

(**A**) Pseudobulked cytokine module (*IFNG*, *TNF*, *IL2*, *IL17A*, *GZMB*, *PRF1*) scores at Wk4 in DPT (left), polyfunctional CD4 T (middle), and polyfunctional CD8 T (right). Lines connect matched unstimulated and WCL-stimulated pseudobulked samples to depict increase in expression following stimulation. *P* values were calculated by paired sign-flip permutations (n=10,000) of pseudobulked module scores.

(**B**) IV specific DPT signature in T cells from an independent cohort. Ridgeline plots of moderated *t* statistics from pseudobulk DE analysis comparing timepoints for less protected (LP) and more protected (MP) macaques for DPT (left), polyfunctional CD4 T (middle), and polyfunctional CD8 T (right) cells from Unstim (top) and WCL-stimulated conditions (bottom). Protection group median is shown by a solid line. Number of samples included in each contrast are shown; differences in counts are due to sample dropout after cell number thresholding.
